# Substrate-derived peptides for selective covalent inhibition of protein tyrosine kinases

**DOI:** 10.64898/2026.05.11.724146

**Authors:** Minhee Lee, Zijing Wang, Andrew C. Johns, Neel H. Shah

## Abstract

Protein tyrosine kinases are important regulators of cell signaling, and aberrant kinase activity contributes to many human diseases, including cancers. All protein tyrosine kinases share a highly-conserved ATP binding pocket but diverge in their substrate binding sites in order to mediate distinct signaling events. Many potent and efficacious ATP-competitive tyrosine kinase inhibitors have been developed, however it remains challenging to achieve on-target selectivity across different kinases and target specific disease mutants, given the high degree of conservation in the ATP-binding pocket. By contrast, the variable substrate-binding site offers an opportunity for selective inhibition, provided molecules can be targeted to this site. Here, we present a modular strategy to design selective, peptide-based covalent inhibitors of tyrosine kinases with a distinct binding mode from existing ATP-competitive inhibitors. Using Src kinase as a model system, we demonstrate that Src-selective reactivity can be achieved by first designing an optimized substrate peptide and then strategically positioning an electrophile on the peptide to target a non-conserved cysteine on the kinase. We show that substrate-derived covalent peptides can inhibit kinase activity, bind simultaneously with an ATP-competitive inhibitor, and even inhibit the activity of kinases bearing a common drug resistance mutation. We further explore the application of this approach to develop an inhibitor of the cancer-relevant fibroblast growth factor receptor 1 kinase that shows selectivity for an oncogenic mutant over the wild-type enzyme. Our modular strategy to generate selective covalent peptides targeting protein tyrosine kinases provides a promising framework for future chemical probe and drug development efforts.

## INTRODUCTION

There are roughly 500 protein kinases encoded in the human genome, 90 of which are classified as protein tyrosine kinases.^1^ Tyrosine kinases drive a broad range of critical biological processes and are central to many cell signaling pathways. As a result, dysregulated kinase activity contributes to many human diseases including cancers and diabetes, making protein tyrosine kinases popular drug targets. Kinase inhibition has traditionally been accomplished by targeting the ATP-binding pocket with small molecules (**Figure 1A**), and there are at least 70 clinically-approved ATP-competitive inhibitors designed to target roughly 20 tyrosine kinases.^2–4^ However, engineering selectivity with ATP-competitive inhibitors is difficult because the ATP-binding site is highly conserved across all protein kinases. In examples where kinase selectivity has been achieved with ATP-competitive inhibitors, the compounds either target or induce unique conformational states in the active site, or are designed with extensive structural analyses of the ATP-binding pocket, exploiting small differences across kinases.^3,5–7^ Furthermore, oncogenic mutations in kinases are often outside of the ATP-binding pocket, making it difficult to design mutant-selective inhibitors. In some cases, selectivity for mutant kinases over their wild-type counterparts has been achieved by exploiting changes in conformational preferences caused by the mutation, or targeting the mutant residue directly during binding.^8–11^ Finally, in cancers, kinase inhibition often leads to drug resistance, and most ATP-competitive inhibitors are notoriously susceptible to mutations within the kinase that disrupt drug binding.^12–15^ Several prior studies have shown that inhibiting a kinase simultaneously with two distinct molecules that bind at different sites can have a synergistic effect and suppress the emergence of drug resistance (**Figure 1B**).^16–19^ However, there is currently no generalizable strategy to identify a druggable second site in kinases that can be systematically applied across the kinome.

**Figure 1.**
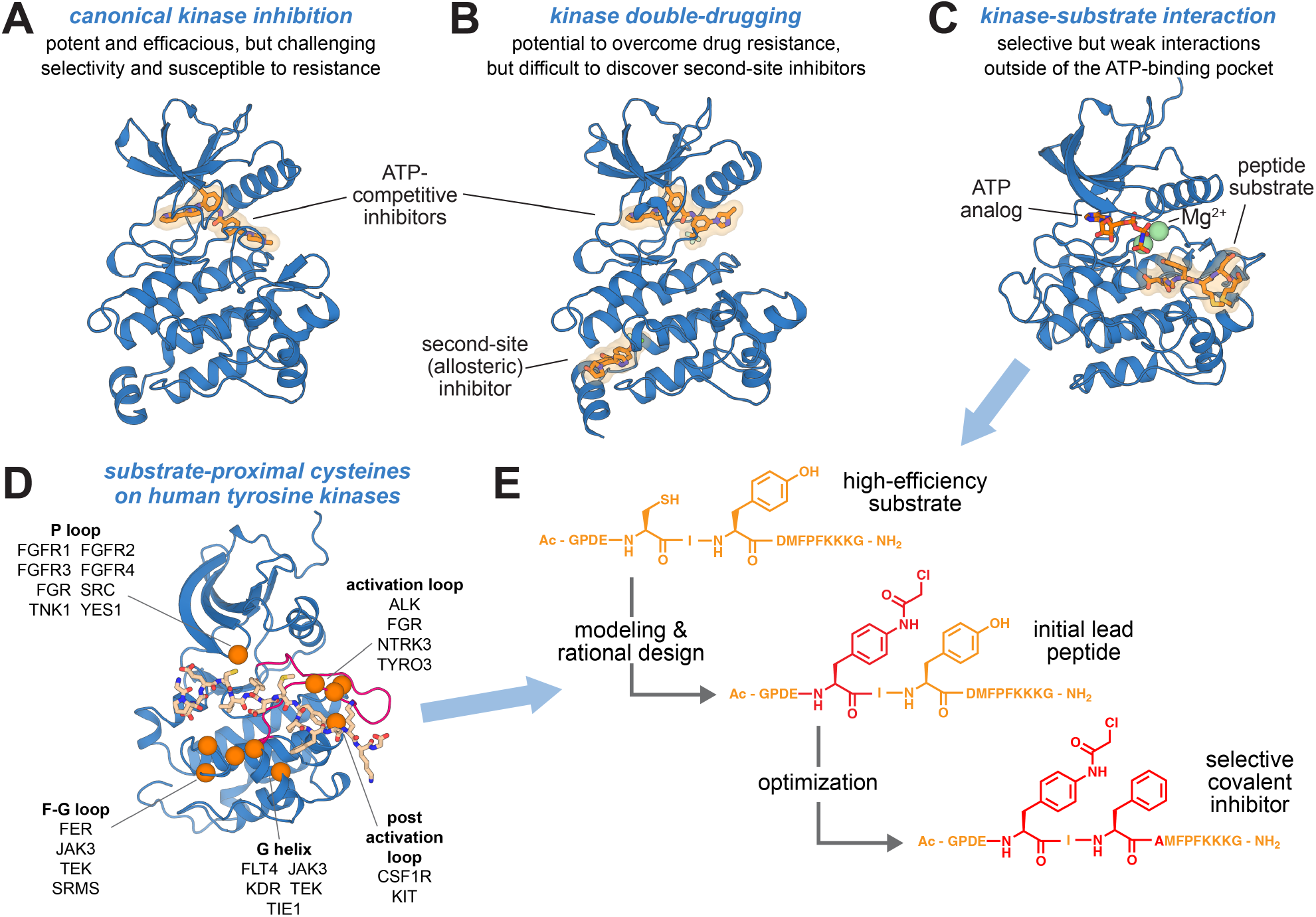
Diverse modes of tyrosine kinase inhibition. (**A**) Canonical inhibition of Abl kinase using the ATP-competitive inhibitor imatinib (PDB code 1OPJ). (**B**) Dual inhibition Abl kinase using the ATP-competitive inhibitor nilotinib and the allosteric inhibitor asciminib (PDB code 5MO4). (**C**) Insulin receptor kinase bound to a peptide substrate (PDB code 1IR3). (**D**) Structural elements on the kinase domain fold where human tyrosine kinases have a cysteine residue near the substrate-binding site, inferred from AlphaFold2 models of active kinases.^40^ See supporting information for details. (**E**) Design of substrate-derived selective covalent kinase inhibitors by exploiting non-conserved cysteine residues and intrinsic sequence specificity.

In this study, we explore the possibility of targeting the substrate-binding region in tyrosine kinases to generate selective inhibitors that are complimentary to ATP-competitive inhibitors (**Figure 1C**). Unlike the highly conserved ATP-binding pocket of kinases, the substrate-binding region is substantially less conserved, as individual kinases have evolved to phosphorylate different target proteins and mediate distinct signaling functions. Several studies have demonstrated that tyrosine kinases recognize specific sequence features surrounding the phospho-acceptor residue on their substrates,^20–22^ and more recently, substrate sequence specificity has been catalogued for nearly every kinase the human kinome.^23,24^ Using substrate-derived peptides or peptidomimetics as selective kinase inhibitors represents a promising and underexploited strategy that complements existing approaches. This strategy has been explored in a number of earlier studies,^25–29^ but potent and selective molecules with this mechanism of action are still lacking for the vast majority of kinases.^30^ This may be partly due to some overt drawbacks of targeting the substrate-binding site, including a relatively shallow pocket (when compared with the ATP-binding pocket) and a lack of conserved molecular scaffolds, other than substrates, themselves, which are generally weak binders (*K*_M_ values in the micromolar range^21^).

We hypothesized that the challenge of weak binding at the substrate-binding site can be addressed by covalent inhibition. Several recent inhibitor design campaigns have exploited covalency to enhance selectivity, target otherwise undruggable sites or proteins, and improve potency or duration of action.^31–34^ Notably, covalent inhibition has been explored extensively for kinase inhibition, but almost exclusively using ATP-competitive molecules.^35–38^ Critically, the substrate-binding sites of many tyrosine kinases have non-conserved cysteine residues (**Figure 1D**). Targeting these cysteines using substrate-derived molecules bearing electrophiles could address the high off-rates of substrates for their kinases and also add a layer of selectivity. Furthermore, although cysteine is the most commonly targeted amino acid, other types of non-conserved residues in the substrate-binding site, including lysine, serine, and threonine, among others, could also conceivably be targeted.^39^

Here, we present a modular strategy to design selective, peptide-based covalent inhibitors of tyrosine kinases with a distinct binding mode from existing ATP-competitive inhibitors (**Figure 1E**). Our approach takes advantage of well-documented kinase substrate specificity profiles and structural models to position electrophiles on substrate-derived peptide scaffolds to react with kinase-specific cysteine residues. We demonstrate that the resulting peptides retain kinase selectivity conferred by the substrate sequence. Critically, selectivity is observed not just across different kinases, but also between a mutant and wild-type kinase that have subtle differences in their substrate preferences. Given their distinct mechanism of action, we also show that our substrate-derived covalent peptides can bind simultaneously with an ATP-competitive small molecule inhibitor. Finally, we demonstrate that these peptides can also inhibit kinases bearing a common mutation that disrupts binding of many ATP-competitive inhibitors. Because the peptides described here are designed using high-throughput substrate profiling data and readily accessible structural models, we envision this pipeline being broadly applicable to most tyrosine kinases. Thus, our study provides a framework for the modular design of kinase-targeted molecules that will complement existing approaches to kinase inhibition.

## RESULTS

### Kinase-selective peptide substrates can serve as scaffolds for covalent ligand design

We recently described a high-throughput bacterial peptide display assay that can be used with custom genetically-encoded peptide libraries to profile substrate specificities of protein tyrosine kinases (**Figure 2A**).^22,41^ Using this approach, we previously screened a degenerate X_5_-Y-X_5_ library with Src kinase domain (Src_KD_), which allowed for the identification of position-specific sequence preferences for Src and the design of a high-efficiency consensus peptide that combined the most enriched residue at each position surrounding the phospho-acceptor tyrosine (**Table 1, peptide 1**).^22^ Notably, in this experiment, we observed strong enrichment of cysteine at the −2 position (**Figure 2B**). Using AlphaFold 3,^42^ we modeled our Src consensus peptide bound to the Src kinase domain and observed that the −2 cysteine on the peptide was close enough to Cys280 on the P-loop of the kinase to potentially form a disulfide bond (**Figure 2C**). To validate this prediction, we incubated the wild-type Src_KD_ with the Src consensus peptide sequence for 1 hour at 37 °C, which revealed a small population of disulfide-adducted Src_KD_ proteins by liquid chromatography-mass spectrometry (LC-MS) (**Figure 2D**). We attribute the low yield of adduct formation to be due to the disulfide bond being reversible in nature, and the reaction taking place in a mildly reducing environment.

**Table 1.** List of covalent peptides used in this study.^a-d^.

| peptide | sequence |
| --- | --- |
| 1 | Ac-GPDE **C** IY DMFPF KKKG-NH<sub>2</sub> |
| 2 | Ac-GPDE **X<sub>1</sub>**IY DMFPF KKKG-NH<sub>2</sub> |
| 3 | Ac-GPDE E IY DMFP**X<sub>1</sub>**KKKG-NH<sub>2</sub> |
| 4 | Ac-GPDE **X<sub>1</sub>**IF DMFPF KKKG-NH<sub>2</sub> |
| 5 | Ac-GPDE **X<sub>1</sub>**I**X<sub>2</sub>**DMFPF KKKG-NH<sub>2</sub> |
| 6 | Ac-GPDE **X<sub>1</sub>**I**X<sub>3</sub>**DMFPF KKKG-NH<sub>2</sub> |
| 7 | Ac-GPDE **X<sub>1</sub>**I**X<sub>4</sub>**DMFPF KKKG-NH<sub>2</sub> |
| 8 | Ac-GPDE **X<sub>1</sub>**I**X<sub>5</sub>**DMFPF KKKG-NH<sub>2</sub> |
| 9 | Ac-GPDE **X<sub>1</sub>**I**X<sub>6</sub>**DMFPF KKKG-NH<sub>2</sub> |
| 10 | Ac-GPDE **X<sub>1</sub>**I**X<sub>7</sub>**DMFPF KKKG-NH<sub>2</sub> |
| 11 | Ac-GPDE **X<sub>1</sub>**I**X<sub>8</sub>**DMFPF KKKG-NH<sub>2</sub> |
| 12 | Ac-GPDE **X<sub>1</sub>**IF AMFPF KKKG-NH<sub>2</sub> |
| 13 | Ac-GPDE **Z<sub>1</sub>**IF AMFPF KKKG-NH<sub>2</sub> |
| 14 | Ac-GPDE **Z<sub>2</sub>**IF AMFPF KKKG-NH<sub>2</sub> |
| 15 | Ac-GPDE **Z<sub>3</sub>**IF AMFPF KKKG-NH<sub>2</sub> |
| 16 | Ac-GPDE **Z<sub>4</sub>**IF AMFPF KKKG-NH<sub>2</sub> |
| 17 | Ac-GPMD **Z<sub>2</sub>**DF WWRPI KKKG-NH<sub>2</sub> |
| 18 | Ac-GPM**Z<sub>2</sub>**G DF WWRPI KKKG-NH<sub>2</sub> |
| 19 | Ac-GPMD **Z<sub>4</sub>**DF WWRPI KKKG-NH<sub>2</sub> |
| 20 | Ac-GPM**Z<sub>4</sub>**G DF WWRPI KKKG-NH<sub>2</sub> |
| 21 | Ac-GPMD **X<sub>1</sub>**DF WWRPI KKKG-NH<sub>2</sub> |
| 22 | Ac-GPM**X<sub>1</sub>**G DF WWRPI KKKG-NH<sub>2</sub> |

a) Non-canonical amino acids X and Z correspond to the following:
X<sub>1</sub> = 4-chloroacetamido-phenylalanine
X<sub>2</sub> = 4-methoxy-phenylalanine
X<sub>3</sub> = 4-acetamido-phenylalanine
X<sub>4</sub> = 4-azido-phenylalanine
X<sub>5</sub> = 4-trifluoromethyl-phenylalanine
X<sub>6</sub> = biphenyl-alanine
X<sub>7</sub> = 4-methyl-phenylalanine
X<sub>8</sub> = 4-amino-phenylalanine

**Figure 2.**
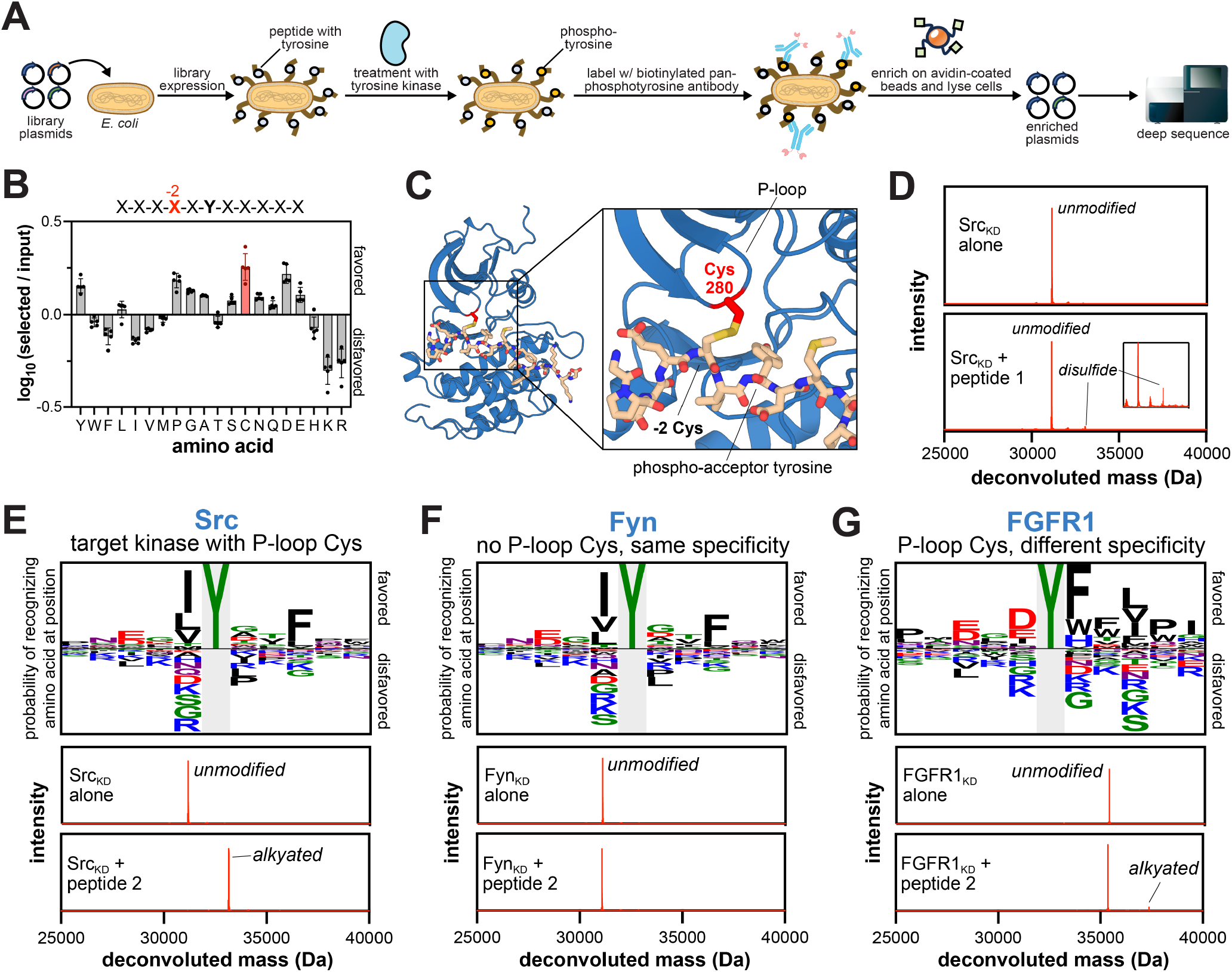
Selective labeling of Src using a substrate-derived peptide. (**A**) Schematic overview of the bacterial display assay method to screen genetically-encoded peptide libraries to identify kinase substrate preferences. (**B**) −2 sequence preferences for Src kinase domain from an X_5_-Y-X_5_ library screen. (**C**) AlphaFold 3 model of the enriched −2 Cys residue in the Src consensus substrate peptide forming a disulfide bridge with the P-loop cysteine (C280) in the active site of Src_KD_. (**D**) Experimental mass spectrometry characterization of Src_KD_ after incubating with 100 μM of the Src consensus **peptide 1** for 1 hour at 37 °C with 2.5 μM Src_KD_. (**E-G**) Probability logos^43,44^ depicting favored and disfavored residues at each position surrounding the phosphotyrosine acceptor, identified from previously reported kinase substrate specificity screens for Src, Fyn, and FGFR1 kinase domains against a ∼10,000 peptide library (*top*).^22^ Each kinase (2.5 μM) was treated with **peptide 2** (500 μM) for 1 hour at 37 °C, revealing distinct selectivity for Src kinase (*bottom*).

To enhance adduct formation and yield an irreversible product, we replaced the −2 cysteine residue on the peptide with a thiol-reactive electrophile, 4-chloroacetamido-phenylalanine (**Table 1, peptide 2**). With this peptide, we observed a significant increase in adducted kinase (**Figure 2E**). Importantly, the observed mass corresponded to a single peptide adduct, despite there being 3 surface-exposed Cys residues on Src_KD_. To confirm that the P-loop cysteine was the reaction site, we made a C280S mutation in Src, which completely abrogated adduct formation (**Figure S2A**). To further validate that the substrate-derived sequence imposes a geometric constraint on chloroacetamide reactivity with the kinase, we moved electrophile to the +5 position (**Table 1, peptide 3**), which is far from the P-loop cysteine in our AlphaFold 3 model. In order to prevent cyclization with the +5 chloroacetamide, the original cysteine residue at the −2 position in the consensus sequence was replaced with a glutamate residue, another preferred amino acid at this position (**Figure 2B**). After incubating **peptide 3** with the WT Src kinase domain, we again observed no adduct formation, in contrast to treatment with **peptide 2** (**Figure S2B**).

Finally, to demonstrate that the peptide sequence taken from kinase specificity profiling assays confers kinase selectivity, we incubated **peptide 2** with two other kinase domains, Fyn and FGFR1, under the same conditions. Fyn kinase has a very similar substrate profile to Src, but it lacks the P-loop cysteine residue. FGFR1 kinase has significantly different substrate specificity profile from Src, but it has a cysteine residue at the same position on the P-loop as Src. Upon incubating **peptide 2** with FGFR1 and Fyn kinase domains, we observed no significant adduct formation for both kinases (**Figure 2F,G**). This suggests that kinase selectivity, as dictated by the primary peptide sequence, is preserved even in the context of a covalently reactive peptide. Importantly, many tyrosine kinases have cysteine residues near the substrate-binding region that are only partly conserved across the kinome (**Figure 1D**). Thus, the additive selectivity from substrate sequence specificity and uniquely positioned cysteine residues could likely be used to design selective covalent peptides for many tyrosine kinases.

### Replacement of the phospho-acceptor residue improves adduct formation

Our initial adduct formation experiments with **peptide 2** were conducted in the absence of ATP. Given the high concentration of ATP in cellular environments, we repeated Src_KD_ labeling in the presence of saturating ATP-Mg^2+^ concentrations. Under these conditions, Src_KD_ can auto-phosphorylate its activation loop, putting it in an active state that should be more competent for substrate binding.^45,46^ However, Src_KD_ could also potentially phosphorylate the central tyrosine residue on **peptide 2**, and the resulting enzymatic product would likely have a fast off-rate that might suppress adduct formation (**Figure 3A**). Indeed, incubating **peptide 2** with Src_KD_ in the presence of ATP-Mg^2+^ revealed a decrease in adduct formation rates (**Figure 3B**).

**Figure 3.**
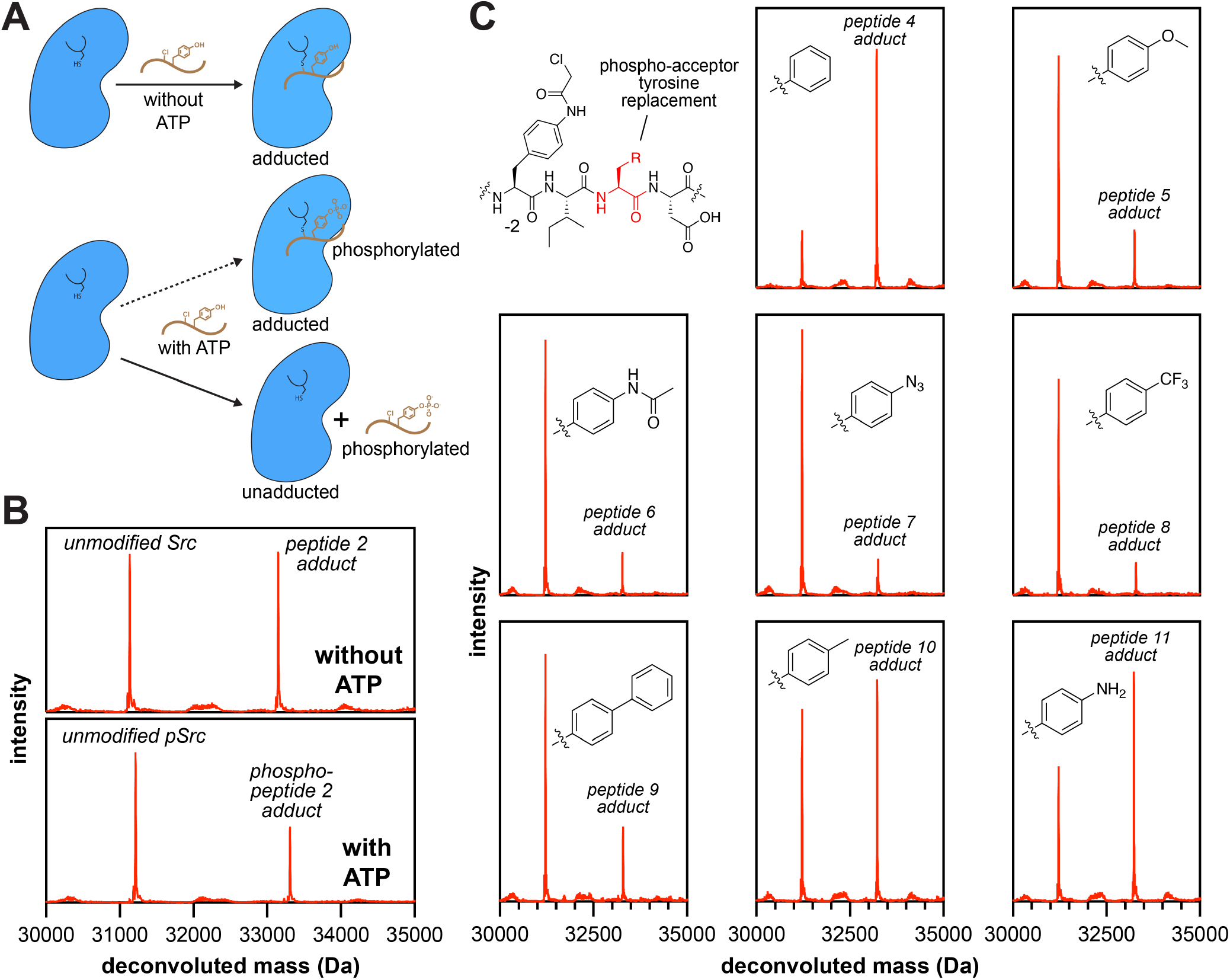
Improving Src covalent labeling by substituting the phospho-acceptor tyrosine. (**A**) Diagram depicting possible outcomes of treating Src with peptide 2 in the absence and presence of ATP. (**B**) MS spectra of **peptide 2** incubated with Src_KD_ (2.5 μM) in the absence or presence of 100 μM ATP for 1 hour at 37 °C. The presence of ATP reduces adduct formation levels. (**C**) MS spectra of tyrosine substitution derivatives of **peptide 2** (**peptides 4-11**, 50 μM) incubated with Src_KD_ (2.5 μM) in the presence of ATP for 1 hour at 37 °C.

To improve adduct formation in the presence of ATP, we investigated various tyrosine and phenylalanine derivatives, with the goal of replacing the tyrosine residue to prevent phosphoryl transfer (**Table 1, peptides 4-11**). We restrained our exploration to tyrosine and phenylalanine derivatives because of highly conserved interactions between the phenyl ring on the substrate tyrosine and residues on the activation loops of tyrosine kinases (**Figure S3A,B**). The modified peptides were incubated with wild-type Src_KD_, and adduct formation was analyzed by LC-MS (**Figure 3C**). Based on prior reports of kinase-substrate interactions, we expected that derivatives which maintained key hydrogen bonding interactions in the kinase active site while preventing phosphoryl transfer (e.g. **peptides 6 and 11** bearing acetamide and amine groups) would demonstrate superior affinity and adduct activity (**Figure S3C**). Surprisingly, however, the Tyr to Phe substitution (**Table 1, peptide 4**) yielded the best performing peptide, suggesting that the phenyl ring is sufficient for binding to the kinase active site (**Figure 3C**).

### Additional structure-activity studies further enhance Src covalent labeling

The consensus sequences derived from high-throughput specificity profiling are generally some of the best peptide substrates identified for tyrosine kinases.^22,47,48^ However, these sequences are derived by assuming that the amino acid preferences at each position around the phospho-acceptor tyrosine are independent of one another. We have previously shown that there is some context dependence in position-specific amino acid preferences.^22,49^ Thus, we looked to improve the primary sequence of the Src consensus sequence to potentially identify tighter binders to the Src active site. We constructed a scanning mutagenesis library based on the Src consensus sequence and screened this library against Src_KD_ using our bacterial peptide display assay (**Figure 1A**) to identify point mutations that enhance activity. From this screen, we identified several point mutations that enhance phosphorylation by Src: D+1A, D+1T, and F+3I (**Figure 4A and Table S3**). These mutant substrates were synthesized, and their activities were orthogonally validated by measuring *in vitro* enzyme kinetics (**Figure S4**). These experiments revealed that the D+1A mutant had the greatest increase in catalytic efficiency, primarily driven by a more than 2-fold tighter *K*_M_ than the base sequence (**Figure S4B**). Based on this result, we introduced the D+1A substitution into **peptide 4** to generate **peptide 12** and compared their ability to label Src_KD_ by LC-MS. As expected, **peptide 12** showed higher adduct formation than **peptide 4**, which was pronounced at lower peptide concentrations, suggesting that the D+1A mutation enhanced binding affinity, mirroring the tighter *K*_M_ seen for the substrate used as a template (**Figure 4B**).

**Figure 4.**
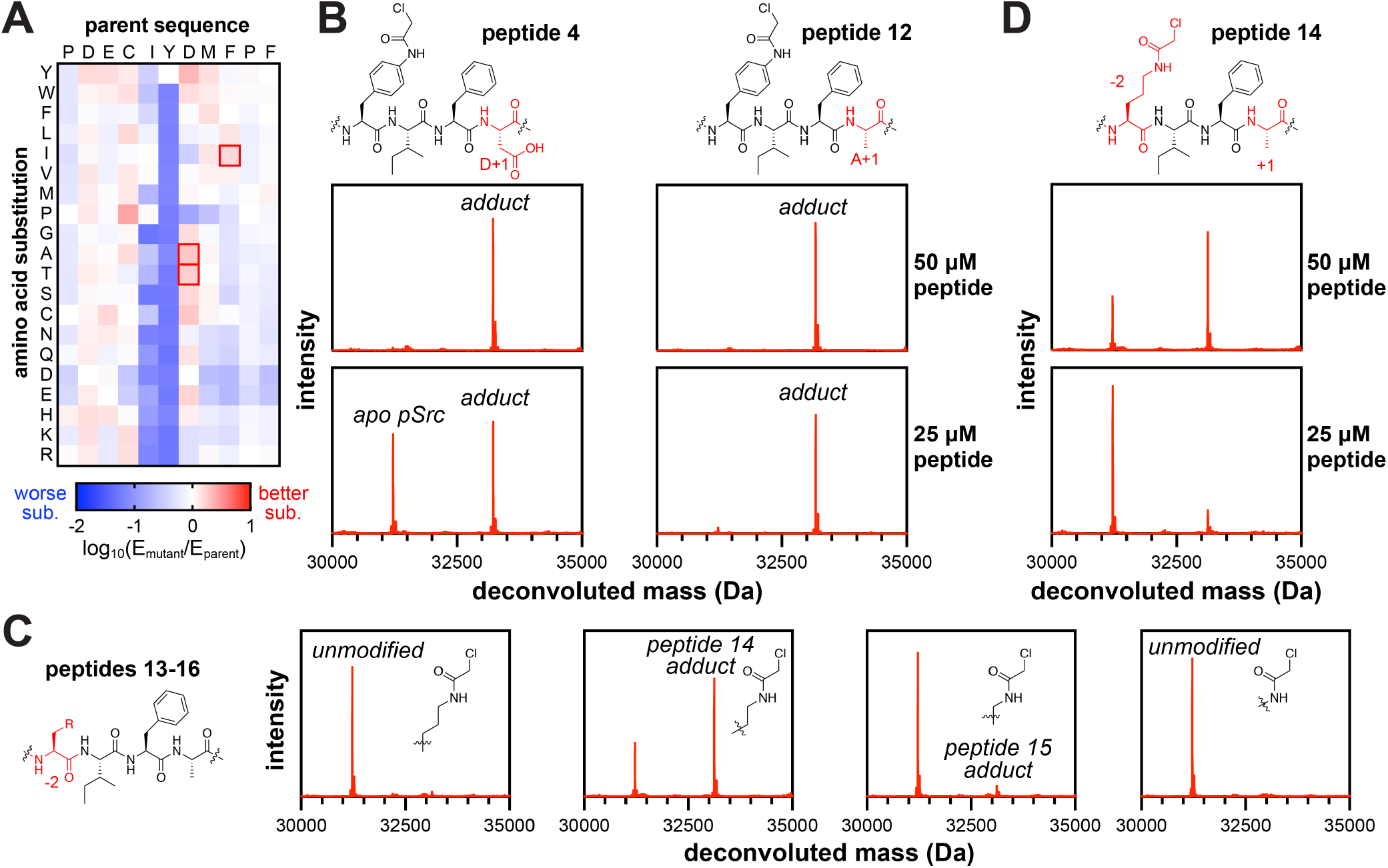
Optimization of the base consensus sequence and electrophile linker in Src-targeting peptides. (**A**) Enrichment matrix of point mutations selected with Src_KD_ from a scanning mutagenesis library of the Src consensus peptide sequence. The top 3 most enriched point mutations (excluding the −2 position and Cys or Tyr substitutions) were identified as D+1A, D+1T, and F+3I. (**B**) MS spectra of **peptide 4** and its best performing point mutant, D+1A (**peptide 12**), both incubated with Src_KD_ (2.5 μM) and either 50 μM or 25 μM peptide at 37 °C for 1 hr in the presence of ATP. The D+1A mutant shows enhanced Src_KD_ labeling at lower concentrations. (**C**) MS spectra of **peptides 13-16** (50 μM), varying alkyl linker length for the −2 chloroacetamide, incubated with Src_KD_ (2.5 μM) at 37 °C for 1 hr in the presence of ATP. Out of the linkers tested, the ornithine derivative (**peptide 14**) provided the best covalent activity. (**D**) MS spectra of **peptide 14** incubated with Src_KD_ (2.5 μM) and either 50 μM or 25 μM peptide at 37 °C for 1 hr in the presence of ATP. Juxtaposition with **peptide 12** in panel **B** shows that the phenyl linker is superior to the ornithine linker.

Lastly, we examined whether the phenyl linker for the chloroacetamide was optimal for adduct formation with the Src_KD_ P-loop. To investigate this, we used alkyl linkers of varying lengths from lysine down to 2,3-diaminopropionic acid (Dap), in the context of the phospho-acceptor Y-to-F and D+1A substitutions (**Figure 4C, peptides 13-16**). The linker variants were then incubated with Src_KD_ and analyzed by LC-MS to observe levels of adduct formation. Of these peptides, only that with the ornithine linker, **peptide 14**, showed appreciable adduct formation (**Figure 4C**). We then compared the ornithine linker to the phenyl linker in the context of the previous two modifications, **peptide 12**, which revealed the phenyl linker still affords more efficient adduct formation (**Figure 4B,D**). Altogether, we identified the combination of the phenyl linker, central phenylalanine, and D+1A mutation as the best performing peptide.

### Src-selective covalent peptides suppress kinase activity

The covalent peptides designed in this study are expected to be substrate-competitive, and thus should inhibit kinase activity. To test this, we first incubated the Src kinase domain with the precursor peptide, **peptide 2**, or the optimized **peptide 12** sufficiently long to ensure complete adduct formation (**Figures 2E and 4B**). Src_KD_ treated with equivalent volume of buffer under the same conditions was included as a non-perturbed control. After complete adduct formation, the kinase/covalent-peptide mixtures were diluted and used to phosphorylate a peptide substrate lacking any cysteine residues (**Figure 5A**). In the absence of a covalent adduct, Src rapidly phosphorylated the substrate (**Figure 5B**). By contrast, the adducted Src_KD_ samples significantly attenuated kinase activity (**Figure 5C,D**). The Src_KD_ adducted with **peptide 2** had more residual activity than Src_KD_ adducted with **peptide 12**. This likely reflects the improved binding of **peptide 12** due to the D+1A mutation, as well as phosphorylation of the phospho-acceptor tyrosine in **peptide 2**, which presumably weakens its occupancy of the active site. To better quantify covalent inhibition parameters, we incubated the Src_KD_ with varying concentrations of **peptide 12** for varying amounts of time, analyzed complex formation by gel shift, and extracted *k*_inact_ (0.8 min^-1^) and *K*_I_ (341 μM) (**Figure S5**). This *K*_I_ was surprisingly weaker than the corresponding *K*_M_ value for the parent substrate peptide (60 μM) (**Figure S4B**), suggesting that substitution of the −2 cysteine for 4-acetamido-phenylalanine and the phospho-acceptor tyrosine for phenylalanine may have an appreciable effect on binding affinity.

**Figure 5.**
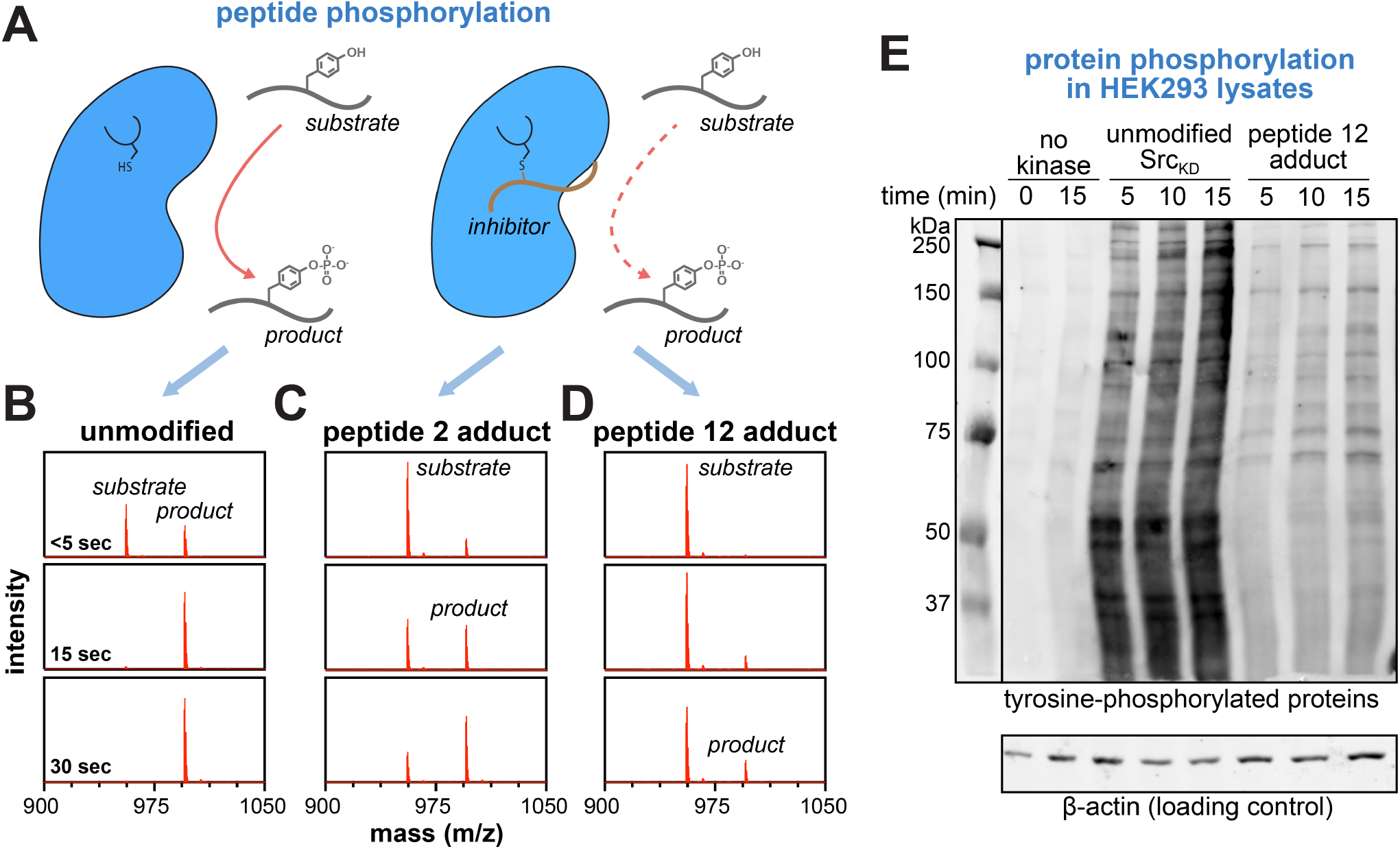
Inhibition of Src kinase activity using covalent peptides. (**A**) Comparison of Src kinase activity with and without adduct formation. (**B**) MS spectra monitoring substrate (Ac-GHDEDIYGMFPFKKKG-NH_2_, 100 μM) phosphorylation by unmodified Src_KD_ (0.5 μM) after <5, 15, and 30 seconds. (**C**) Same as panel **B**, but with Src_KD_ fully adducted by **peptide 2**. (**D**) Same as panels **B** and **C**, but with Src_KD_ fully adducted by **peptide 12**. (**E**) Phospho-tyrosine blot of HEK293 lysates treated with ATP-Mg^2+^ and either no kinase, unmodified Src_KD_, or Src_KD_ fully adducted by **peptide 12**, showing significantly reduced phosphorylation of intact proteins by the adducted kinase.

We next tested if Src_KD_ modification by our substrate-derived peptides impacted the ability of Src to phosphorylate intact proteins, rather than short peptides. We used HEK293 cell lysates as a source of diverse protein substrates, which we phosphorylated with apo Src_KD_ as a non-perturbed control or Src_KD_ adducted with **peptide 12**. Phosphoprotein levels were analyzed by western blotting with a pan-phosphotyrosine antibody, revealing significant attenuation of protein phosphorylation by adducted Src_KD_ (**Figure 5E**). This result confirms that our covalent peptides can also outcompete intact protein substrates to inhibit kinase activity.

It is noteworthy that the fully adducted Src_KD_ proteins still exhibited some residual activity against both peptide and protein substrates. We hypothesize that this is likely due to incomplete anchoring of the covalently attached peptide in the substrate-binding pocket, allowing substrates to still access the active site. This could be the result of the P-loop-attached peptide rotating away from the active site. Alternatively, it is noteworthy that solution structural studies and simulations support the idea that there are two, mostly non-overlapping binding modes for substrate peptides on tyrosine kinases.^50^ It is plausible that our covalent peptides only preclude one of these binding modes. To explore this idea, we generated an AlphaFold 3 model of Src_KD_ including both Src consensus **peptide 1** and a second cysteine-less variant of the same peptide (**Figure S6**). Remarkably, this model reveals an arrangement for two peptides bound to one kinase domain, with one covalently attached to the P-loop cysteine via a disulfide bond. While this is only a prediction and lacks supporting experimental evidence, it suggests a plausible explanation for partial inhibition by our covalently-attached substrate-derived peptides. Critically, the structure-activity relationship studies described in the preceding sections suggest that systematic improvements can be made to these covalent peptides to suppress kinase activity, and further modifications could yield even more efficacious inhibitors than the ones presented here.

### Covalent peptides are complementary to ATP-competitive small molecules

One potential application of substrate-competitive inhibitors could be to “double-drug” a kinase, in conjunction with established ATP-competitive inhibitors. To explore this possibility, we examined if our covalent Src inhibitors can bind to Src_KD_ simultaneously with dasatinib, a well-established ATP-competitive inhibitor that binds Src and some other kinases with sub-nanomolar affinity.^51^ Dasatinib is a type I kinase inhibitor that binds deep in the ATP-binding pocket, and it binds Src in a conformation that should be compatible with substrate binding (**Figure 6A**). To test whether our covalent peptides and dasatinib could engage Src simultaneously, Src_KD_ was pre-incubated with saturating concentrations of dasatinib, then treated with **peptide 12**. Covalent adduct formation was monitored by gel-shift analysis (**Figure 6B**). Rapid peptide adduct formation was observed in both cases, with a 1.5-fold attenuation of the rate in the presence of dasatinib (**Figure 6C**). We suspect that this slight reduction in reaction kinetics may be due to previously reported mild negative cooperativity between the ATP- and substrate-binding sites.^52^

**Figure 6.**
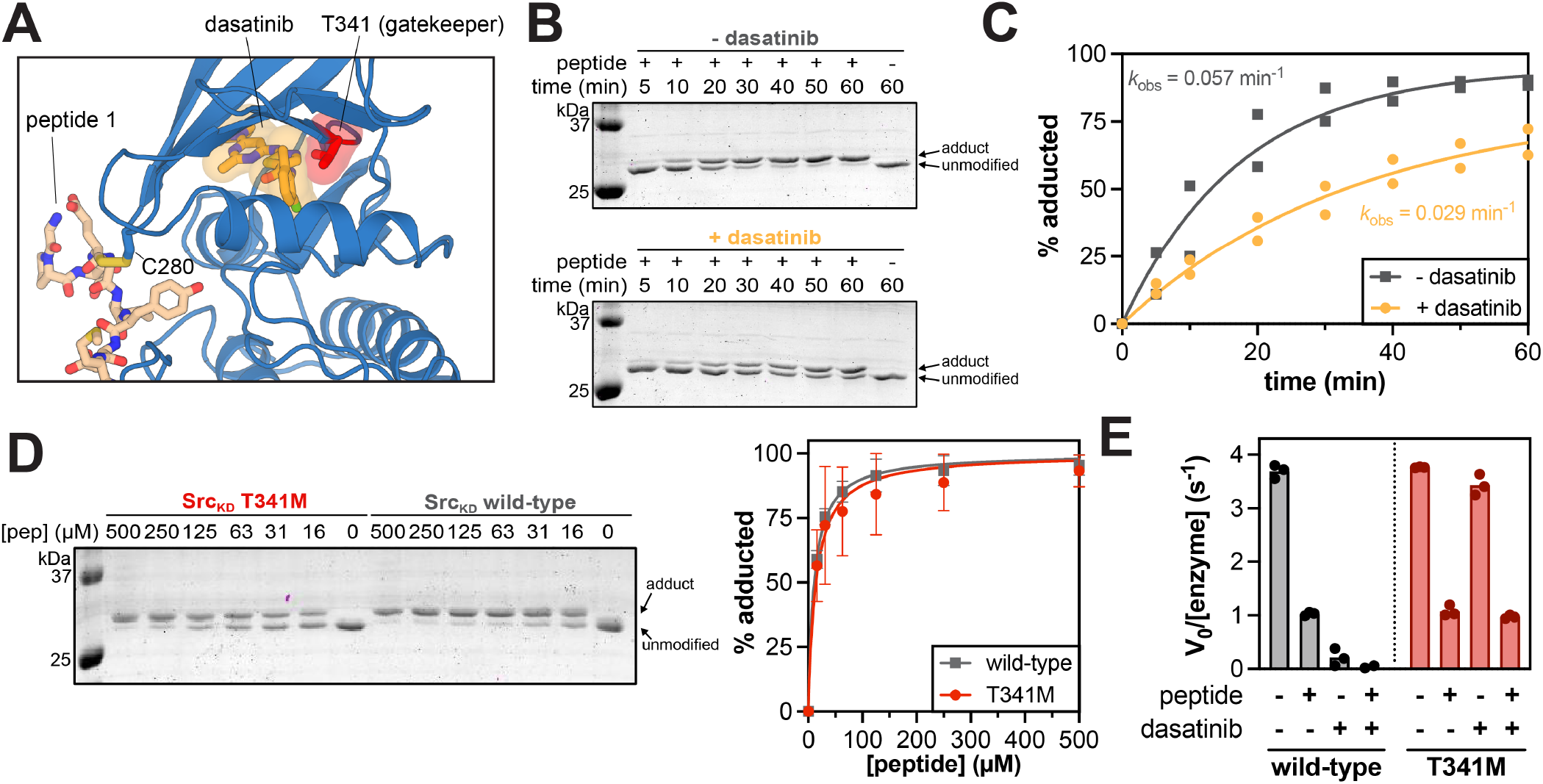
Covalent peptides are compatible with and bind orthogonally to ATP-competitive inhibitors. (**A**) AlphaFold 3 model of Src bound to **peptide 1** (same as **Figure 2C**), with the positioning dasatinib (orange) shown, based on superposition of the AlphaFold 3 model with a Src-dasatinib co-crystal structure (PDB code 3G5D). The gatekeeper residue, T341, is highlighted in red. (**B**) Gel-shift assay of Src_KD_ (2.5 μM) pre-incubated with DMSO or 10 μM dasatinib, followed by 63 μM of **peptide 12** at 37 °C. (**C**) Quantified gel-shift assay data in (B) from two replicates. (**D**) Adduct formation with Src_KD_ wild-type or gatekeeper mutant T341M (2.5 μM) and **peptide 12** at varying peptide concentrations after 30 minutes of incubation at 37 °C. Adduct formation was monitored by gel-shift (left) and three replicate experiments were quantified (right). (**E**) Kinase activity of Src_KD_ WT and Src_KD_ T341M against a substrate peptide (Ac-GHDEDIYGMFPFKKKG-NH_2_, 63 μM) when pre-treated with 250 μM **peptide 12**, 50 nM dasatinib, or both, compared to the untreated Src_KD_ (three replicates).

Next, we hypothesized that, since **peptide 12** binds orthogonally to ATP-competitive compounds, it may still inhibit some Src mutants that perturb dasatinib binding. One of the most common resistance mutations to ATP-competitive kinase inhibitors is at the gatekeeper residue, where typically a smaller amino acid (e.g. threonine) is substituted for a bulkier one (e.g. methionine or isoleucine) (**Figure 6A**).^53^ We first confirmed that **peptide 12** was still able to form a covalent adduct with Src_KD_ in the presence of the clinically-observed Src gatekeeper mutation, T341M (**Figure 6D**). Next, we examined whether dasatinib and **peptide 12** could inhibit Src_KD_ wild-type and Src_KD_ T341M, alone and in conjunction. The activity of Src_KD_ variants was then measured against a substrate peptide in the presence and absence of dasatinib, with and without a **peptide 12** adduct (**Figure 6E**). As expected, 50 nM dasatinib almost completely inhibited Src_KD_ wild-type but did not inhibit Src_KD_ T341M. However, when treated with saturating concentrations of **peptide 12**, both the wild-type protein and the T341M mutant had suppressed kinase activity, confirming that the T341M mutation confers dasatinib resistance but does not impact covalent peptide binding and inhibition. Finally, we note that for Src_KD_ wild-type, combination of both dasatinib and **peptide 12** was additive, leading to full suppression of kinase activity (**Figure 6E**). Altogether, our data demonstrate that our covalent peptide inhibitor binds to Src in a conformation that is compatible with type I ATP-competitive inhibitors, suggesting that this binding mode is potentially useful for “double-drugging” kinases. Furthermore, the distinct mechanism of kinase engagement by these peptides may be useful in the face of common resistance mutations.

### Substrate-derived covalent peptides enable mutant-specific kinase inhibition

Some prevalent disease mutations in tyrosine kinases lie in or around the substrate-binding region, raising the possibility that our substrate-templated covalent peptides could be designed to be mutant-selective. As a noteworthy example, there are oncogenic mutations in fibroblast growth factor receptor (FGFR) kinases that drive a broad range of cancers,^54,55^ and one of the most prevalent such mutations in FGFR1, FGFR2, and FGFR3 is a Lys to Glu charge reversal mutation near the active site (**Figure 7A and Figure S7**).^56^ This mutation is reported to enhance kinase activity by stabilizing an active conformation of the kinase,^57^ but we hypothesized that it could also impact substrate specificity. To study potential changes in substrate specificity profiles, we screened wild-type FGFR1_KD_ and the K656E mutant against a library containing ∼10,000 proteome-derived sequences (10K library)^22^ using our bacterial display platform (**Figure 7B and Table S4**). Although there was a strong correlation in peptide preferences between the wild-type and K656E mutant kinases, we observed a selective phosphorylation of a subset of the library by the mutant kinase, suggestive of a change in substrate specificity. This change in specificity was further validated using *in vitro* enzyme kinetics measurements using purified peptides derived from the screen (**Figure S8**). A deeper examination of the preferred substrate sequence features for each FGFR1 variant revealed that the charge reversal correlated with a complementary change in electrostatic preferences, particularly at the +2 and +3 residues, where basic residues such as +2R and +3R went from disfavored by FGFR1 wild-type to neutral or favored by FGFR1 K656E, and acidic residues such as +2D and +3D were less favorable for the mutant kinase over wild-type (**Figure 7C**). These substrate residues are expected to be in close proximity to the mutation site (**Figure 7A**).

**Figure 7.**
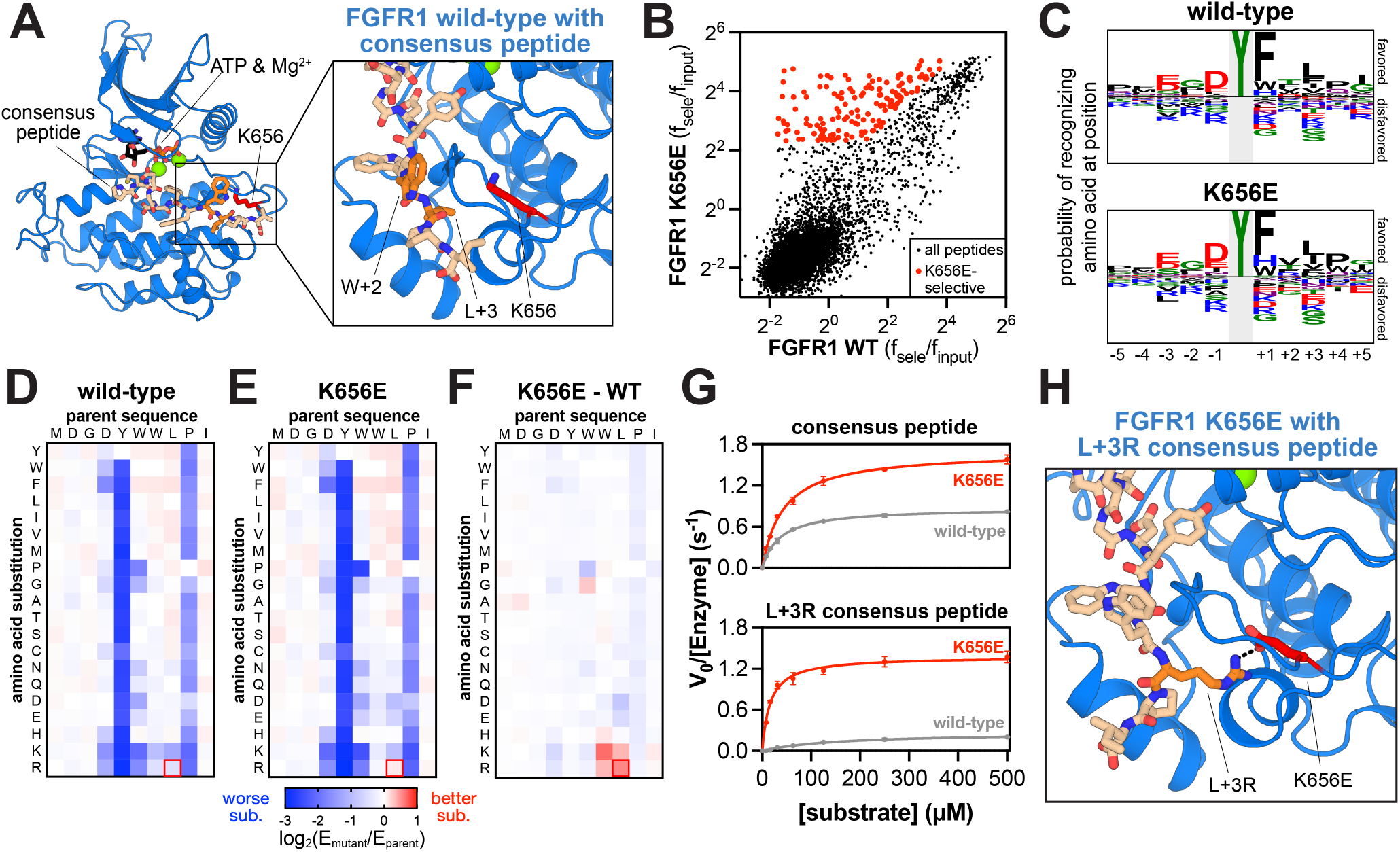
A common oncogenic mutation in FGFR kinases alters substrate preferences. (**A**) AlphaFold 3 structure model of FGFR1_KD_ bound to the FGFR1 consensus substrate peptide. The K656 residue points towards the +2 and +3 residues of the peptide substrate, which may impact the residue preferences at these sites with a Lys to Glu mutation. (**B**) Scatterplot of enrichment values following sorting of the 10K library with FGFR1_KD_ WT and K656E. Data shown are the average of three independent replicates. A population of mutant-selective sequences is highlighted. (**C**) Probability logos derived from the most enriched peptides for each kinase, based on the data shown in panel **B**. (**D**) and (**E**) Scanning mutagenesis matrices highlighting the effects of substitutions on the FGFR1 consensus base sequence for phosphorylation by the wild-type and K656E kinase domains (average of three replicates). Enrichments for each mutant are expressed on a log_10_ scale and normalized to the enrichment of base sequence (set to zero). (**F**) Difference between the matrices in panels (D) and (E), highlighting that K/R are favored by K656E over wild-type at the +2 and +3 positions. (**G**) Michaelis-Menten curves of the base FGFR1 consensus peptide and the L+3R consensus peptide against FGFR1_KD_ wild-type and K656E variants (three replicates). (**H**) AlphaFold 3 model of the K656E mutant kinase bound to the L+3R consensus sequence.

Although the proteome-derived peptide library screen revealed clear changes in substrate specificity, this analysis did not yield a very selective high-efficiency substrate. As a complementary approach, we conducted a scanning mutagenesis bacterial display screen starting from a high-efficiency FGFR1 consensus peptide, comparing FGFR1_KD_ wild-type and K656E (**Figure 7D,E and Table S5**). Although both FGFR1 variants showed very similar sensitivity to mutations in the consensus substrate (e.g. intolerance to mutations at the +4 Pro residue), subtraction of the two mutational scanning matrices revealed a stark preference for basic residues at the +2 and +3 position by FGFR1 K656E over wild-type (**Figure 7F**). Enzyme kinetic measurements revealed that, whereas both FGFR1 wild-type and K656E phosphorylated the original FGFR1 consensus peptide with similarly high catalytic efficiencies, the L+3R mutant peptide was a markedly better substrate for the mutant kinase (**Figure 7G and Figure S8E**). Finally, AlphaFold 3 modeling of the L+3R peptide with FGFR1 K656E revealed a direct ion pair between Glu656 on the mutant kinase and Arg+3 on the substrate providing a clear structural explanation for this enhancement in activity and selectivity (**Figure 7H**).

Having identified a selective substrate sequence for the K656E mutant, we next looked to apply the same design principles as discussed earlier to generate a selective, covalent peptide inhibitor for the FGFR1 K656E mutant. As noted earlier, FGFR kinases have a cysteine on the P-loop, like Src, suggesting that similar positioning of the electrophile might be appropriate (**Figure 1D**). However, upon modeling the FGFR1 consensus substrate peptide in the active site of FGFR1_KD_ K656E with AlphaFold 3, we noticed that the N-terminal half of the peptide adopts an α-helical conformation, and thus the −2 position may not be well-positioned for an electrophile to react with the P-loop cysteine (**Figure 8A**). Prior studies with EGFR substrates corroborate this notion, in which the identity of the −1 residue (hydrophobic, as in the Src consensus, versus acidic, as in the FGFR1 consensus) can dictate the secondary structure of the N-terminal half of substrate peptides when docked on a kinase domain.^49^ Thus, we tested a series of peptides containing chloroacetamides at the −2 and −3 positions, with three different linkers (**Table 1, peptides 17-22**). With an ornithine or Dap linker (**peptides 17-20**), we observed an enhancement in covalent labeling of FGFR1 with a peptide containing a −3 electrophile over a −2 electrophile (**Figure 8B and Figure S9**). This positional preference was inverted with the phenyl linker (**peptides 21-22**), however we also observed a second FGFR1 adduct with these peptides at later time points, suggesting some non-specific labeling (**Figure S9**). Aryl chloroacetamides are generally more intrinsically reactive than alkyl chloroacetamides,^58^ and while this was not an issue for the Src-targeting molecules, we reasoned that this may be a liability in the context of the FGFR1-targeting peptide. Thus, we opted to proceed with a peptide bearing a chloroacetamide attached via −3 Dap linker (**peptide 20**). Finally, we confirmed that **peptide 20** was a selective covalent modifier of the FGFR1 K656E mutant over the wild-type kinase (**Figure 8C**).

**Figure 8.**
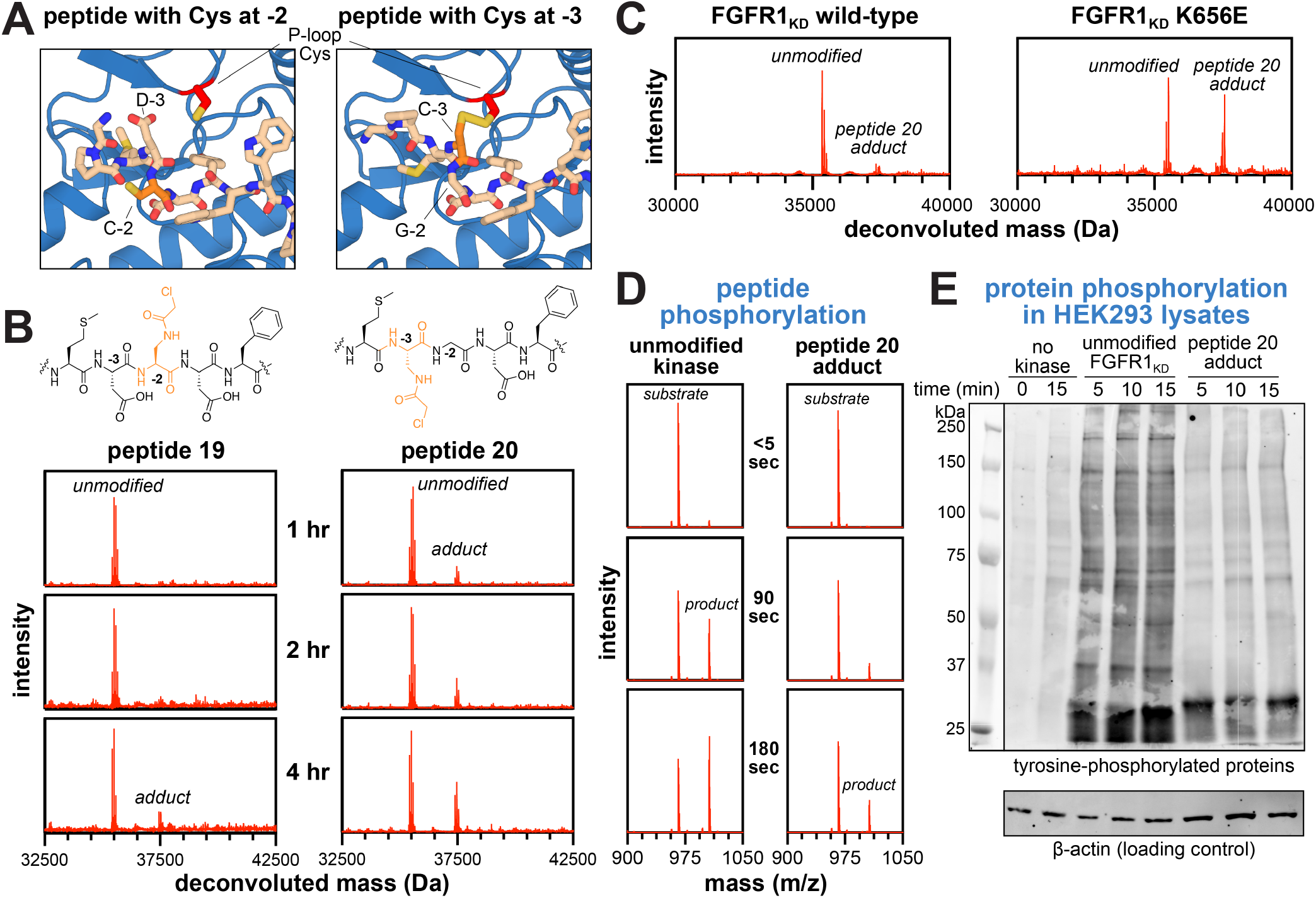
Conversion of an FGFR1-mutant selective substrate into a covalent inhibitor. (**A**) AlphaFold 3 models of FGFR1 consensus peptide substrate. The −2 position was predicted to be out of position for reactivity with the P-loop cysteine, while the −3 position was in better position. The electrophile was installed at the −3 residue position. (**B**) MS data showing that adduct formation with FGFR1_KD_ K656E (2.5 μM) is better with the chloroacetamide at the −3 position (**peptide 20**, 100 μM) than the −2 position (**peptide 19**, 100 μM). Note that FGFR1 is pre-activated by auto-phosphorylation, resulting in an ensemble of phosphorylation-states (**Table S2**).^62^ (**C**) MS spectra of FGFR1 WT and FGFR1_KD_ K656E (2.5 μM) incubated with 100 μM **peptide 20** for 4 hours at 37 °C, showing significant mutant selectivity. (**D**) Phosphorylation of a substrate peptide (Ac-GPMDGDYWWLPIKKKG-NH_2_, 100 μM) by FGFR1_KD_ K656E (1 μM) unadducted or pre-incubated with **peptide 20** (100 μM) for 4 hours at 37 °C. (**E**) Phospho-tyrosine blot of HEK293 lysates treated with ATP-Mg^2+^ and either no kinase, unmodified activated FGFR1_KD_ K656E, or activated FGFR1_KD_ K656E adducted by **peptide 20**, showing significantly reduced phosphorylation of intact proteins by the adducted kinase.

With the optimized, selective covalent peptide for FGFR1 K656E (**Table 1, peptide 20**), we next analyzed kinase inhibition against peptide and protein substrates. We observed significant attenuation of FGFR1_KD_ K656E kinase activity toward a peptide substrate when adducted by **peptide 20** (**Figure 8D**). We also treated HEK293 lysates with FGFR1_KD_ K656E and confirmed that adduct formation suppresses kinase activity against intact protein substrates (**Figure 8E**). Taken together with our Src-targeting peptides, these results demonstrate the generalizability of our design strategy. Furthermore, it is noteworthy that our strategy could be used to produce an FGFR mutant-selective molecule. While there have been some recent successes in designing ATP-competitive inhibitors that are selective for individual FGFR kinase members (e.g. FGFR3 over FGFR1/2/4), including some that are showing promising efficacy in the clinic, these molecules do not differentiate between wild-type and mutant kinases.^59–61^

## DISCUSSION

Here we report a strategy to design selective, covalent peptide inhibitors of protein tyrosine kinases. This strategy combines our extensive understanding of tyrosine kinase substrate specificity with structural models to pinpoint substrate proximal cysteine residues for covalent targeting. Using this approach, we demonstrated that a covalent peptide based on the Src consensus sequence can site-specifically react with Src and inhibit its kinase activity. We also demonstrated that the unique binding mode of these peptides means that they can bind a kinase simultaneously with some ATP-competitive inhibitors, and they are still efficacious in the presence of a common drug resistance mutation that disrupts binding of many ATP-competitive inhibitors. A critical feature of our method is that we can achieve a high degree of selectivity by combining substrate sequence preferences with carefully positioned electrophiles that target non-conserved cysteine residues. For example, our Src-targeting molecules do not label Fyn kinase, whereas very few ATP-competitive molecules can differentiate between these two highly-homologous enzymes.^63^ Similarly, guided by substrate specificity differences, we designed a peptide that selectively targets the FGFR1 K656E cancer mutant over the wild-type enzyme, which has not been reported for other FGFR-family inhibitors.^59–61^

The results described above illustrate the modularity of our method, however our approach has several limitations that warrant discussion. For example, while we were able to take advantage of changes in substrate specificity in the context of FGFR1 K656E, it is noteworthy that only mutations that perturb the substrate-binding site can be exploited using our approach. We acknowledge that not all mutations will have this effect, in which case alternative strategies will be required to selectively target those specific mutants. On the other hand, although we focused on a mutation that is directly involved in substrate recognition, one could also envision using our approach to target kinase mutants with allosteric effects on substrate binding.^64^ Another limitation to our approach, which is common to all covalent inhibitors, is the emergence of resistance mutations at the targeted nucleophilic residues.^65^ For instance, for both Src and FGFR1 kinases, mutations at the P-loop cysteine would ablate inhibition by our covalent peptides. Furthermore, it is noteworthy that the P-loop cysteine in Src appears to be important in regulating kinase activity through redox mechanisms.^66,67^ Transient oxidation of this cysteine would also disrupt covalent targeting by canonical cysteine-reactive functional groups, underscoring the need to target other non-conserved side chains or even specific redox states of cysteine.^68,39^ Finally, we acknowledge that kinases adducted by our substrate-derived peptides still retain partial activity. While inhibition could be improved through further optimization of the peptides, one could also envision exploiting this residual activity in the context of an induced proximity modality, whereby the kinase is selectively recruited to a target protein to induce a neo-phosphorylation event that modulates signaling.^69^

Overall, this study illustrates the opportunities and benefits afforded by covalently targeting the substrate-binding site in tyrosine kinases. We recognize that the structure-activity relationship studies shown here for our proof-of-concept peptides are not exhaustive. Further improvements, aided by computational methods and more exhaustive structure-activity studies will likely yield more potent and selective covalent inhibitors than the ones reported here. For instance, our scanning mutagenesis-based optimization was constrained to single substitutions on the existing consensus sequences, but different starting points or higher-order mutations may yield more potent molecules. Additionally, we note that the molecules reported here are exclusively linear peptides, which have notable challenges in metabolic stability and membrane permeability as a therapeutic.^70^ Peptide cyclization has been a popular approach to address these challenges, improving binding affinity, proteolytic stability, bioavailability, and membrane permeability.^70^ Many selection methods involving phage and mRNA display to directly screen cyclic peptides have been reported.^71,72^ Substrate-competitive peptidomimetics have also been screened against target kinases, expanding the chemical diversity of potential peptide-based inhibitor compounds.^73^ Leveraging these methods to covalently target the substrate-binding site with chemically diverse molecules could yield inhibitors with enhanced potency and bioavailability, while preserving the selectivity and orthogonality to ATP-competitive inhibition, as illustrated in our study.

## ASSOCIATED CONTENT

### Supporting Information

The Supporting Information PDF file contains a detailed Materials and Methods section and Supplementary Figures S1-S9. The Supporting Information spreadsheet file contains Tables S1-S5.

### Data availability

All data needed to evaluate the results presented here are provided in the main text and figures or in the Supplementary Materials. AlphaFold 3 models of kinase-peptide complexes are available via the Columbia University Academic Commons (https://doi.org/10.7916/74ey-2j83).

## Author contributions

The manuscript was written through contributions of all authors. All authors have given approval to the final version of the manuscript. Specific contributions are given below:

Conceptualization: M.L., N.H.S.

Data curation: M.L., N.H.S.

Formal analysis: M.L., N.H.S.

Funding acquisition: N.H.S.

Investigation: M.L., Z.W., A.C.J.

Methodology: M.L., A.C.J., N.H.S.

Project administration: N.H.S.

Resources: N.H.S.

Supervision: N.H.S.

Validation: M.L.

Visualization: M.L., N.H.S.

Writing – original draft: M.L., N.H.S.

Writing – review & editing: M.L., Z.W., A.C.J, N.H.S.

## Notes

Columbia University has filed a patent application related to this work for which N.H.S. and M.L. are inventors.

## ACKNOWLEDGEMENT

We thank members of the Shah lab for their insightful discussions, and for their technical and conceptual guidance throughout this project. This work used resources from the Columbia University Department of Chemistry Mass Spectrometry Core Facility, the Precision Biomolecular Characterization Facility (PBCF), and the JP Sulzberger Columbia Genome Center. We thank Fereshteh Zandkarimi for her assistance with mass spectrometry, and we thank Jerry Chang and Martin Acosta for training and support in the PBCF.

