## Supplementary Figures and Methods for "Substrate-derived peptides for selective covalent inhibition of protein tyrosine kinases"


### Table of contents


Materials and methods pg 2
Figure S1. RP-HPLC analysis of all covalent peptides used in this study. pg 7
Figure S2. Covalent peptides target a specific Cys residue in the Src P-loop. pg 8
Figure S3. Key interactions at the phospho-acceptor tyrosine in kinase active sites. pg 9
Figure S4. Michaelis-Menten analysis of Src consensus point mutants. pg 10
Figure S5. Determination of $k_{inact}$ and $K_I$ parameters for Src with peptide 12. pg 10
Figure S6. AlphaFold 3 model of two peptides bound in the Src active site. pg 10
Figure S7. The landscape of cancer mutations in FGFR kinases. pg 11
Figure S8. Validation of changes in FGFR1 substrate specificity caused by the K656E mutation. pg 12
Figure S9. Analysis of electrophile linker and position for FGFR K656E targeting. pg 13
Supplementary references pg 13


**Supplementary tables included as separate spreadsheet file**

Table S1. Mass spectrometry analysis of covalent peptides used in this study.
Table S2. Expected and observed unmodified and adducted protein masses.
Table S3. Phosphorylation of the Src consensus scanning mutagenesis library by Src kinase.
Table S4. Phosphorylation of 10K library by FGFR1 wild-type and K656E.
Table S5. Phosphorylation of the FGFR1 consensus scanning mutagenesis library by FGFR1 wild-type and K656E.

### MATERIALS AND METHODS

#### *Peptide synthesis and purification*

All peptides were synthesized using 9-fluorenylmethoxycarbonyl (Fmoc) solid-phase peptide chemistry. All syntheses were carried out using the Liberty Blue automated microwave-assisted peptide synthesizer from CEM under nitrogen atmosphere, with standard manufacturer-recommended protocols. Peptides were synthesized on MBHA Rink amide ProTide resin (0.1 mmol scale). The following Nα-amino acids were used in the peptide syntheses described in this paper: all 20 canonical Nα-Fmoc-L-amino acids with standard side-chain protecting groups (CEM), Nα-Fmoc-L-(Alloc)lysine-OH (Ambeed), Nα-Fmoc-L-(Alloc)ornithine-OH (Ambeed), Nα-Fmoc-L-(Alloc)-2,4-diaminobutyric acid (Dab)-OH (Ambeed), Nα-Fmoc-L-(Alloc)-2,3-diaminopropionic acid (Dap)-OH (Ambeed), Nα-Fmoc-(4-nitro)-L-phenylalanine-OH (TCI America), Nα-Fmoc-(4-methoxy)-L-phenylalanine-OH (Alfa Aesar), Nα-Fmoc-(4-acetylamino)-L-phenylalanine-OH (Sigma), Nα-Fmoc-(4-azido)-L-phenylalanine-OH (ChemImpex), Nα-Fmoc-(4-trifluoromethyl)-L-phenylalanine-OH (Thermo Scientific), Nα-Fmoc-L-4,4'-biphenylalanine-OH (Ambeed), Nα-Fmoc-(4-methyl)-L-phenylalanine-OH (Thermo Scientific), Nα-Fmoc-(4-amino)-L-phenylalanine-OH (Ambeed).

Each Nα-Fmoc-amino acid (6 eq, 0.2 M) was activated with diisopropylcarbodiimide (DIC, 1.0 M) and ethyl cyano(hydroxyamino)acetate (OxymaPure, 1.0 M) in N,N-dimethylformamide (DMF) prior to coupling. Each coupling cycle was done at 75 °C for 15 s followed by 90 °C for 110 s. Deprotection of the Fmoc group was performed in 20% (v/v) piperidine in DMF (75 °C for 15 s then 90 °C for 50 s). The resin was washed 4x with DMF following Fmoc deprotection and after Nα-Fmoc amino acid coupling. All peptides were acetylated at the N-terminus with 10% acetic anhydride/DMF and washed 4x with DMF after the acetylation reaction.

Following peptide synthesis, the resin was washed with DMF, dichloromethane (DCM), and methanol (MeOH) and dried under reduced pressure overnight. The peptides were cleaved from resin and the side chains were simultaneously deprotected in 95% (v/v) trifluoroacetic acid (TFA), 2.5% (v/v) water, and 2.5% (v/v) triisopropylsilane (TIPS), in a ratio of 10 µL of cleavage cocktail per mg of resin. The cleavage-resin mixture was incubated at room temperature for 90 min, with agitation. The cleaved peptides were precipitated in cold diethyl ether, pelleted, and dried under air. The peptides were dissolved in 50% (v/v) acetonitrile/water solution and filtered from resin. The filtrate was freeze-dried for downstream purification.

The crude peptide mixture was purified using reverse-phase high performance liquid chromatography (RP-HPLC) on a preparatory C18 column (XBridge Peptide BEH C18, OBD Prep Column, 130Å, 19x150mm, 5µm) and Waters system. Flow rate was maintained at 17 mL/min with solvents A (water, 0.1% (v/v) TFA) and B (acetonitrile, 0.1% (v/v) TFA). Peptides were generally purified over a 23 min linear gradient from 0-70% solvent B. Peptide purity was assessed using an analytical column (Agilent, ZORBAX 300 SB-C18, 4.6x150mm, 5µm) at a flow rate of 1 mL/min over a 5-95% solvent B gradient in 10 min on an Agilent system. All peptides were determined to be ≥95% pure by peak integration. The identities of the peptides were confirmed using mass spectrometry (Waters Xevo G2-XS QTOF). Pure peptides were lyophilized and redissolved in Tris buffer (100 mM, pH 8.0) or DMSO, as needed for experiments. Concentrations of peptides with canonical amino acids and alkyl chloroacetamides were determined by measuring absorbance at 280 nm and using well-established extinction coefficients for the aromatic amino acids. Concentrations of peptides with phenyl chloroacetamides and other phenylalanine derivatives were estimated by analytical HPLC quantification, measuring absorbance at 214 nm, as previously reported.[1] Phenylalanine derivatives were assumed to contribute to 214 nm absorbance to a similar extent as phenylalanine.

#### *Synthesis of N-chloroacetamide electrophiles on peptides*

All chloroacetamide couplings were done with chloroacetyl chloride (Alfa Aesar). Peptides containing chloroacetamide warheads were synthesized in one of two ways depending on the attached linker. Prior to chloroacetamide coupling, all peptides were synthesized as described with the Liberty Blue automated microwave-assisted peptide synthesizer under nitrogen atmosphere and using standard manufacturer-recommended protocols.

Peptides with the chloroacetamide warhead on a phenyl linker were first synthesized with Nα-Fmoc-(4-nitro)-L-phenylalanine-OH, incorporated where the electrophile is desired along the peptide sequence. The nitro group was then reduced to the amine,[2] then coupled with chloroacetyl chloride, as follows: After synthesis of the full peptide sequence bearing 4-nitrophenylalanine, the resin was swelled in 3 mL of dimethylformamide (DMF) for 45 min at room temperature (RT) in a scintillation vial. Tetrahydroxydiboron (0.9 mmol, 9 equiv.) was added to the stirring suspension. A 0.75 mM solution of 4,4'-bipyridine in DMF was separately prepared in a scintillation vial. 2 mL of the 4,4'-bipyridine solution (0.15 mol % relative to peptide on resin) was added to the stirring resin mixture and left at RT for 10 min. After generation of 4-aminophenylalanine, the solvent was removed and the resin was washed in 5 mL DMF. The resin was resuspended in 5 mL DMF and potassium carbonate (2.2 mmol, 22 equiv.) was added. On ice, chloroacetyl chloride (2.1 mmol, 21 equiv.) was added dropwise to the stirring mixture. The reaction was brought up to RT, stirring for 1 hr. The mixture was then washed with water and resin was filtered from solvent through a fritted column. The resin was washed DMF, dichloromethane (DCM), and methanol (MeOH) and dried under reduced pressure overnight. The peptides were then deprotected and cleaved as described above.

Peptides with the chloroacetamide warhead on an alkyl linker were synthesized with Alloc-protected versions of the amino acid, such as Nα-Fmoc-(Alloc)-L-lysine-OH, incorporated where the electrophile is desired along the peptide sequence. Following the synthesis of the full peptide sequence, the resin was swelled in 5 mL DMF for 45 min at RT in a scintillation vial. The DMF was removed and the resin was resuspended in 5 mL DCM. Phenylsilane (40 mmol, 400 equiv.) was added to the stirring mixture. The Alloc protecting group was removed with tetrakis(triphenylphosphine)palladium(0) (52 mmol, 520 equiv.) which was added to the suspension and stirred at RT for 4 hrs, protected from light. The resin was decolorized with a 20 mL solution of diethyldithiocarbamate (0.67 mmol, 6.7 equiv.) and diisopropylethylamine (0.57 mmol, 5.7 equiv.) dissolved in DMF. The resin was washed with DMF, resuspended in 5 mL DMF, then transferred to a scintillation vial. Potassium carbonate (2.2 mmol, 22 equiv.) was added to the mixture. On ice, chloroacetyl chloride (2.1 mmol, 21 equiv.) was added dropwise to the stirring mixture. The reaction was brought up to RT, stirring for 1 hr. The mixture was then washed with water and resin was filtered from solvent through a fritted column. The resin was washed DMF, dichloromethane (DCM), and methanol (MeOH) and dried under reduced pressure overnight. The peptides were then deprotected and cleaved as described above.

#### *Plasmid mutagenesis*

All wild-type protein kinase expression vectors were reported previously.[3] The kinase domain mutants were cloned using Agilent QuickChange primer design. Primers ligating to the ends of the kinase domain gene were designed and paired with the QuickChange primers to clone the kinase domain gene into two fragments containing the desired point mutation. The fragmented kinase domain gene was re-ligated via PCR using the insert primers. The reconstituted kinase domain gene insert was ligated into the pET23a vector backbone by Gibson assembly. The plasmids were sequenced via whole plasmid sequencing using Plasmidsaurus to confirm the presence of the mutation change.

#### *Expression and purification of tyrosine kinase domains*

Constructs for the kinase domains all contained an N-terminal $His_6$-tag followed by a TEV protease cleavage site, and were encoded in the pET23a expression vector (AmpR). Kinase domains were co-expressed in *E.coli* BL21(DE3) cells with the YopH tyrosine phosphatase (StrepR). Cells transformed with the kinase domain and YopH were grown in Terrific Broth (LB) supplemented with 100 μg/mL ampicillin and 100 μg/mL streptomycin at 37°C. Once cells reached an optical density of 0.5-0.6 at 600 nm, 0.5 mM isopropyl-β-D-1-thiogalactopyranoside (IPTG) was added to the cultures to induce protein expression and were incubated at 18°C overnight (16-18h). Cells were harvested by centrifugation at 4000xg at 4°C for 30 min and resuspended in lysis buffer (50 mM Tris, pH 7.5, 300 mM NaCl, 20 mM imidazole, 2 mM β-mercaptoethanol, 10% glycerol) with protease inhibitor cocktail [200 μM AEBSF (Calbiochem), 20 μM leupeptin (Calbiochem), 1 μM peptstatin A (Sigma)], and lysed by sonication. Lysates were separated from insoluble material by centrifugation (33,000xg at 4°C for 45 min) and the supernatant was filtered through 1.1 μm syringe filters. The filtered supernatant was purified over a 5 mL HisTrap Ni-NTA column (Cytiva), followed by anion exchange on a 5 mL HiTrap Q column (Cytiva), and eluted with a gradient of 50 mM NaCl to 1 M NaCl in 50 mM Tris, pH 8.5, 1 mM TCEP-HCl, and 10% glycerol. $His_6$-TEV tags were cleaved with 0.1 mg/mL TEV protease for 2 hrs at room temperature. Cleavage mixtures were separated over Ni-NTA columns and concentrated by centrifugation in Amicon Ultra-15 30 kDa MWCO spin filters (Millipore). The concentrate was separated on a Superdex 75 16/600 gel filtration column (Cytiva), equilibrated with 10 mM HEPES, pH 7.5, 100 mM NaCl, 1 mM TCEP-HCl, 5 mM $MgCl_2$, 10% glycerol. Collected fractions were validated by SDS-PAGE, pooled, and concentrated in the Amicon spin filters. Purified proteins were aliquoted and stored at -80°C. Concentrations of purified proteins were determined by measuring absorbance at 280 nm and using well-established extinction coefficients for the aromatic amino acids.

#### *Pre-activation of FGFR1 kinase domains*

Fibroblast growth factor receptor-1 (FGFR1) kinase domains were pre-activated prior to use in biochemical assays and experiments. 25 μM FGFR1 kinase domains were incubated with 5 mM adenosine triphosphate (ATP) for 1 hour at 37°C. The phosphorylated kinase domains were re-concentrated in 10 mM HEPES, pH 7.5, 100 mM NaCl, 1 mM TCEP-HCl, 5 mM $MgCl_2$, and 10% glycerol, and concentration was re-verified by Nanodrop. The pre-phosphorylated FGFR1 kinase domain constructs were stored at -80°C.

#### *Bacterial peptide display assay*

Electrocompetent MC1061 cells (Lucigen) were transformed with genetically-encoded bacterial display peptide libraries and screened as described previously.[3,4] A detailed step-by-step protocol for preparing the libraries, displaying the peptides, phosphorylating libraries with a purified kinase, enriching phosphorylated cells, deep sequencing the samples, and analyzing data can be found in reference 2.

#### *Visualizing kinase specificity data as probability logos*

The probability logos shown in **Figure 2** and **Figure 7** are derived from peptide display screens using the 10K phosphosite library (also known as the pTyr-Var Library).[3] Enrichment scores for each peptide in the library by taking the frequency of a sequence in the phosphorylated and selected sample over its frequency in the input library. Enrichments from three independent replicates were averaged. For all logos, only sequences bearing a central tyrosine were analyzed (7382 sequences). For the Src, Fyn, and FGFR1 data in **Figure 2**, from previously reported screens,[3] a cutoff enrichment score of 2 was used (Src = 727 sequences, Fyn = 594 sequences, FGFR1 = 363 sequences). For the new FGFR1 wild-type and K656E screens reported here (**Figure 7**), a cutoff of 5 was used (wild-type = 154 sequences, K656E = 170 sequences). The log odds of the binomial probability of observing each feature was calculated using the pLogo webserver,[5] with sequences above the cutoff as the "foreground" and the full list of 7382 sequences as the "background". Numerical probability values were exported from the pLogo webserver, then plotted using the Python Logomaker package.[6] The height of the central tyrosine in each logo is arbitrarily scaled to the maximum stack size in the positive direction.

#### *Identification of substrate-proximal Cys residues across human tyrosine kinases*

To identify cysteine residues that are near the substrate-binding region (**Figure 1D**), we analyzed previously reported AlphaFold2 models of every tyrosine kinase in an active conformation.[7] All models were aligned to a structure of the insulin receptor kinase bound to a substrate (PDB code 1IR3), then visually inspected for nearby cysteine residues. Kinases containing cysteine residues near the substrate were grouped based on the structural element containing the cysteine, as shown in the figure.

#### *Mass spectrometry analysis of kinase adduct formation with covalent peptides*

Covalent adduct formation between kinase domains and peptides was detected by LC-MS (Waters UPLC H-Class system coupled to Xevo G2-XS Q-ToF). Covalent adduct reactions were done with 2.5 µM kinase domain treated with 100 µM ATP and varying concentrations of the covalent peptide of interest. The reactions were quenched with β-mercaptoethanol. Samples were then injected onto a Waters LC column (ACQUITY Premier Protein BEH C4 300Å, 1.7 µm, 2.1x100mm). An initial flow rate of 0.425 mL/min was maintained for the first 1.25 min with solvents A (water + 0.1% formic acid) and B (acetonitrile + 0.1% formic acid), then reduced to 0.300 mL/min for 4.75 min, for a total run time of 6 min. The samples were separated over a linear gradient of 5% to 85% B over 2.5 min, followed by a 95% B wash step. The multi-charge state spectra from electrospray ionization for each sample were deconvoluted using the Maximum Entropy (MaxEnt) algorithm.

#### *Gel-shift analysis of kinase adduct formation with covalent peptides*

Covalent adduct reactions with kinase domains were done with 2.5 µM kinase domain treated with 100 µM ATP and varying concentrations of the covalent peptide of interest. Time points or end points were quenched with 4X sample buffer, then boiled for 5 min. Samples were separated on 7.5% precast polyacrylamide gels via SDS-PAGE. Gels were stained with Coomassie Blue [0.1% Coomassie R-250 (w/v), 10% acetic acid (v/v), 40% ethanol (v/v)] and destained with destaining solution [10% ethanol (v/v), 7.5% acetic acid (v/v)]. Gels were imaged with the Amersham Typhoon 5 biomolecular imager (Cytiva) on the short IR fluorescent setting, and quantified using ImageJ software.

#### *Kinase activity assays measuring peptide phosphorylation*

Kinase activities were measured using the ADP Quest assay by Eurofins, following standard protocols provided by the manufacturer. All experiments reported here were set up in 384 well-plates. Peptide solutions were diluted in 100 mM Tris, pH 8.0 to desired concentrations. The final reaction mixtures contained 10 nM kinase domain with 100 µM ATP. Kinase reactions were initiated by adding 1 mM ATP into a 50 µL reaction mixture for a final concentration of 100 µM ATP. Phosphorylation reaction progress was monitored by tracking fluorescence at excitation 530 nm and emission 590 nm every 90 sec at 37°C on a plate reader (BioTek Synergy Neo2). The fluorescence units (RFU) were converted to µM ADP by standard curve.

Michaelis-Menten analyses for peptide substrates of tyrosine kinases were done by extracting the initial rates of phosphorylation reaction from the linear regimes of the reaction progress curves. Initial rates were also taken from negative control samples containing a kinase but lacking the peptide substrate, to account for background ATP hydrolysis. This background rate was subtracted from the rates measured with peptide substrate present. The adjusted initial rates were plotted as a function of peptide substrate concentration and fit to the Michaelis-Menten equation to determine the $k_{cat}$ *and* $K_M$ kinetic parameters.

For measuring the activities for kinases treated with covalent peptides and dasatinib, 2.5 µM of the tyrosine kinase domains were first pre-treated with 250 µM of covalent peptide and 100 µM ATP, diluted in 50 mM Tris, pH 7.5, 150 mM NaCl, 5 mM $MgCl_2$ for 30 min at 37°C. A comparative apo control was prepared alongside the peptide-treated sample, instead adding equivalent volume of 100 mM Tris, pH 8.0 to the reaction mixture. The final reaction mixtures for this experiment contained 10 nM kinase domain (treated or untreated with covalent peptide), 63 µM substrate, 50 nM dasatinib, and 100 µM ATP. Kinase phosphorylation reactions were initiated by adding 630 µM substrate into a 50 µL reaction mixture for a final concentration of 63 µM substrate. The initial rates for each sample were extracted from the linear regimes of the reaction progress curves. Initial rates were also taken from negative control samples containing the corresponding kinase domain treatment conditions but lacking the peptide substrate to account for variability in background ATP hydrolysis. This background rate was subtracted from the initial rates corresponding to the matching sample conditions. The adjusted initial rates were then compared across treatment conditions.

#### *Cell culture and transfection*

Cells were cultured in a 37°C tissue culture incubator with 5% $CO_2$. Cells were discarded by passage 25 and tested for mycoplasma every 6 months. HEK293 cells were grown in Dulbecco's Modified Eagle Medium (DMEM) with 10% Fetal Bovine Serum (FBS) and 1% penicillin/streptomycin.

3 x $10^6$ HEK293 cells were seeded in 10 cm well plates and transfected 24 hours later with 10 µg DNA in 1 mL empty DMEM using 30 µL of polyethylenimine (PEI). The next morning the medium was replaced with full DMEM. 24 hours later each plate was harvested in PBS by scraping and spun down at 1000xg for 5 minutes. Cell pellets were flash frozen at -80°C until further processing. The cell pellets were then thawed and lysed in 450 µL of lysis buffer (20 mM Tris-HCl, pH 8.0, 137 mM NaCl, 10% glycerol, and 1% Triton X-100 + protease inhibitors, no EDTA or phosphatase inhibitors) for 25 minutes on ice. The cell lysate mix was then spun down at 17,000xg for 15 minutes and supernatant transferred to new Eppendorf tubes and stored at -20°C. Protein concentration was determined with a bicinchoninic acid (BCA) assay and absorbance was measured at 562 nm using a plate reader (BioTek Synergy Neo2).

#### *Kinase activity assays measuring protein phosphorylation in lysates*

Kinase domain constructs were prepared with and without covalent peptide for comparative analysis. 5 µM of kinase domain were treated with 100 µM ATP and 250 µM covalent peptide, and incubated at 37°C for 30 min. The negative control samples were treated with Tris buffer in place of covalent peptide.

100 µL of lysate were transferred to new Eppendorf tubes and $MgCl_2$, ATP, and kinase reaction mixtures were added to final concentrations of 5 mM, 2 mM, and 1 µM respectively. Lysates were then incubated at 37°C for 15 minutes with aliquots taken out at 0-, 5-, 10-, and 15-minute time points. 15 µg of protein lysate was loaded onto a 12% polyacrylamide gel and separated via SDS-PAGE. Gels were transferred onto a nitrocellulose membrane using TurboBlot (BioRad) and the membrane was blocked with 5% Bovine Serum Albumin (BSA) in Tris-buffered saline (TBS) for 1 hour at room temperature. Membranes were rinsed with TBS with 0.1% Tween-20 (TBS-T) and incubated with primary antibodies in TBS-T + 5% BSA overnight at 4°C. The following primary antibodies were used, β-Actin (Cell Signaling Technologies, #4970S), 1:1000 and Phosphotyrosine (Sigma Aldrich, #05-321X) 1:1000. Membranes were washed 5x with TBS-T and incubated with secondary antibodies (IRDye® 680RD Goat anti-Rabbit IgG, LiCor, #926-68071; and IRDye® 800CW Goat anti-Mouse IgG, LiCor, #926-32210). The blots were then washed again 5x with TBS-T and imaged on a Cytiva (Amersham) Typhoon 5 imager.

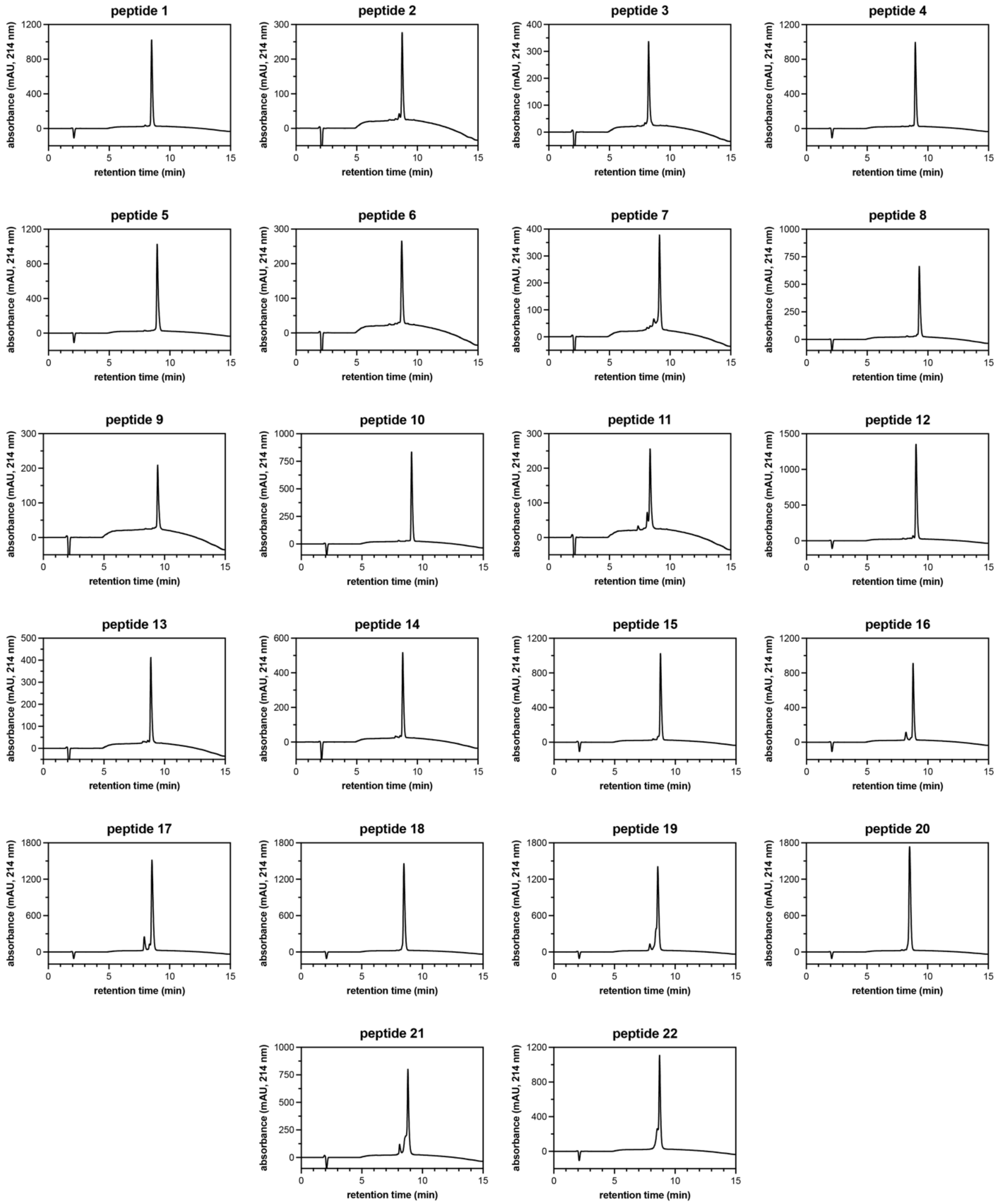


**Figure S1. RP-HPLC analysis of all covalent peptides used in this study.** All peptides were run on an Agilent ZORBAX 300 SB-C18, 4.6x150 mm, 5 µm analytical HPLC column at a flow rate of 1 mL/min. Solvent A was water with 0.1% trifluoroacetic acid, and Solvent B was acetonitrile with 0.1% acetic acid. Samples were analyzed using the following program: initial 2 minute isocratic phase in 5% B, then a linear 10 minute gradient from 5-95% B over 10 minutes, followed by a hold at 95% B.

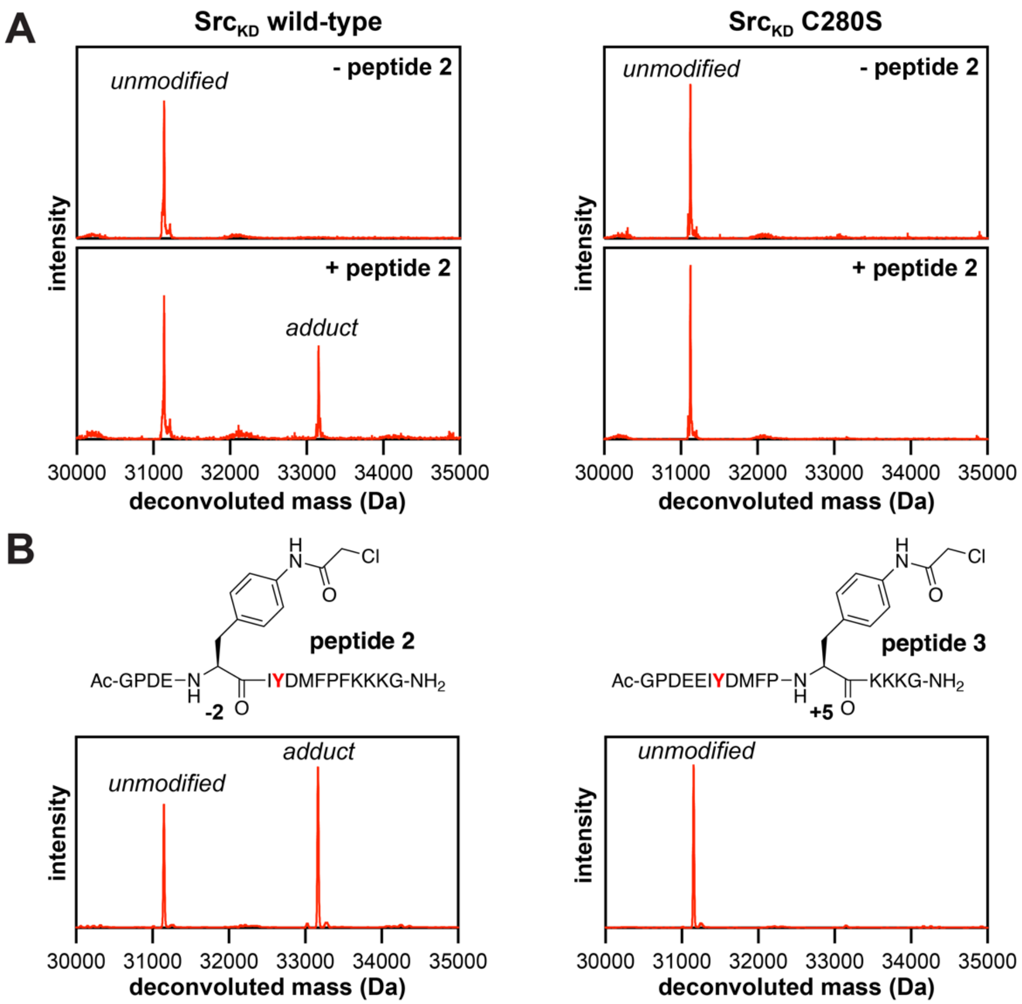


**Figure S2. Covalent peptides target a specific Cys residue in the Src P-loop.** (**A**) MS spectra of **peptide 2** (50 μM) incubated with Src$_{KD}$ wild-type or C280S (2.5 μM) for 1 hour at 37 °C. Loss of adduct formation with the C280S mutation confirms that P-loop C280 is the main adduct site. (**B**) MS spectra of **peptide 2** and **peptide 3** (50 μM) incubated with Src$_{KD}$ wild-type or C280S (2.5 μM) for 1 hour at 37 °C. Only **peptide 2** with the electrophile correctly positioned for reaction with the P-loop cysteine has adduct formation, indicating that adduct formation is also dependent on correct positioning of the electrophile on the peptide sequence.

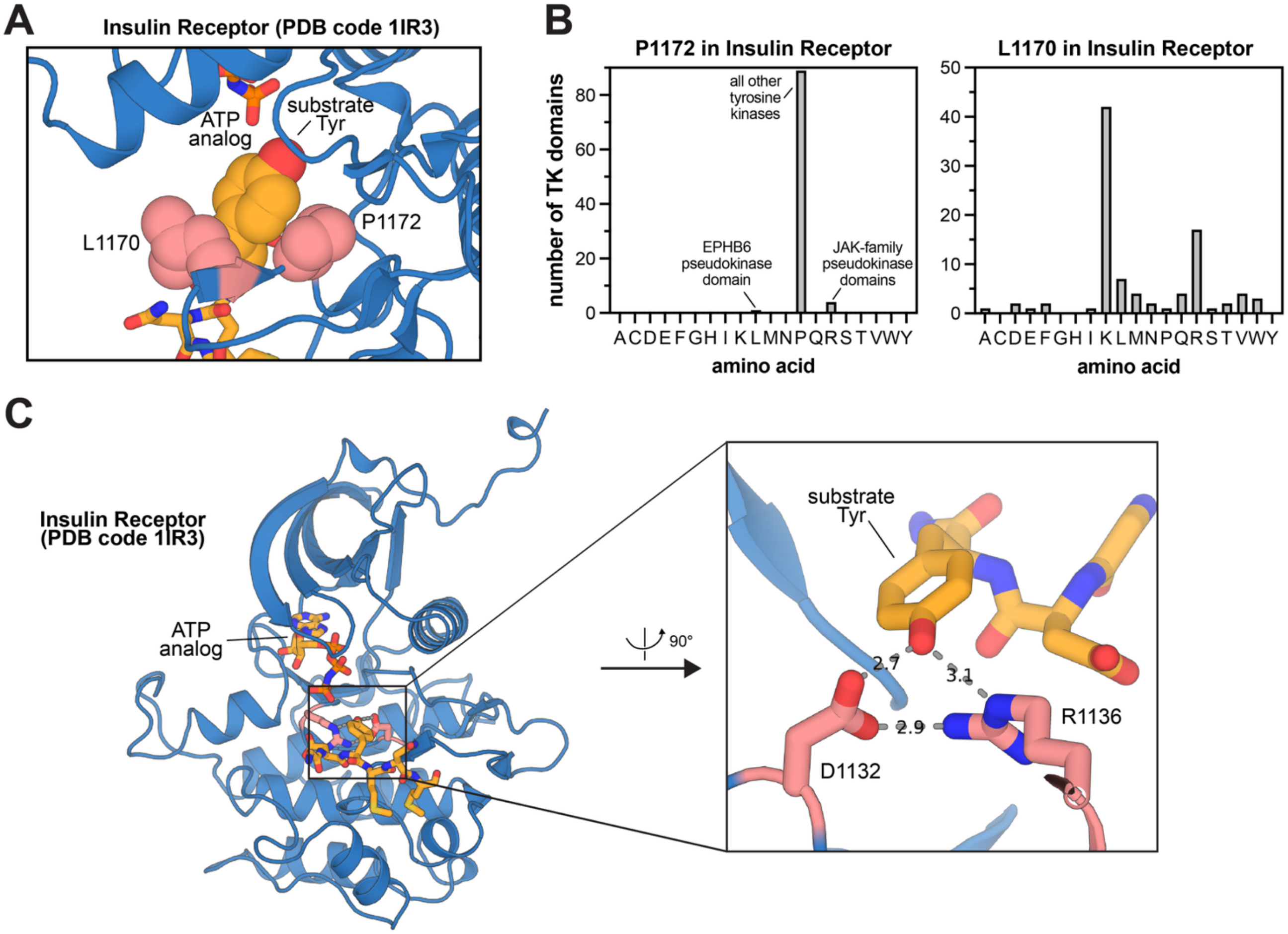


**Figure S3. Key interactions at the phospho-acceptor tyrosine in kinase active sites.** (**A**) Tyrosine phenyl ring interactions. CH/π interactions with an invariant proline (P1172 in Insulin Receptor kinase PDB code 1IR3, shown, and P428 in Src). Nearly invariant hydrophobic or cation/π interactions two residues before the proline (L1170 in Insulin Receptor kinase PDB code 1IR3, shown, and K426 in Src). (**B**) Conservation of key residues that interact with the phospho-acceptor tyrosine phenyl ring. (**C**) Key H-bonding interactions at the phospho-acceptor residue with an aspartate and arginine on the highly conserved catalytic loop of tyrosine kinases (PDB code 1IR3).

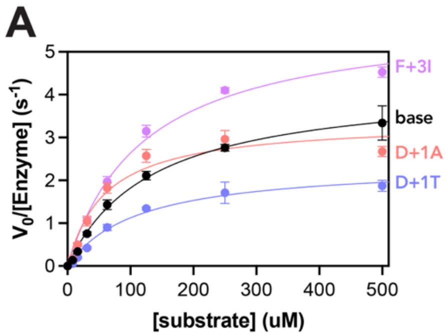


B

| name | sequence | $k_{cat}$ ($s^{-1}$) | $K_M$ (μM) | $k_{cat}/K_M$ $(M*s)^{-1}$ |
|---|---|---|---|---|
| consensus (peptide 1) | Ac-GPDECIYDMFPFKKKG-NH2 | 4.33 ± 0.51 | 140 ± 33 | 31,959 ± 4,182 |
| consensus, D+1A | Ac-GPDECIY**A**MFPFKKKG-NH2 | 3.38 ± 0.13 | 60 ± 3 | 56,795 ± 4,193 |
| consensus, D+1T | Ac-GPDECIY**T**MFPFKKKG-NH2 | 2.40 ± 0.21 | 113 ± 9 | 21,278 ± 928 |
| consensus, F+3I | Ac-GPDECIYDM**I**PFKKKG-NH2 | 5.85 ± 0.20 | 122 ± 10 | 48,112 ± 2,141 |

**Figure S4. Michaelis-Menten analysis of Src consensus point mutants.** (**A**) Michaelis-Menten analysis plot of the point mutants for the Src consensus **peptide 1**. (**B**) Kinetic parameters for each peptide as shown in (A). The D+1A mutant demonstrated the lowest $K_M$ value, indicative of tighter binding to the kinase active site.

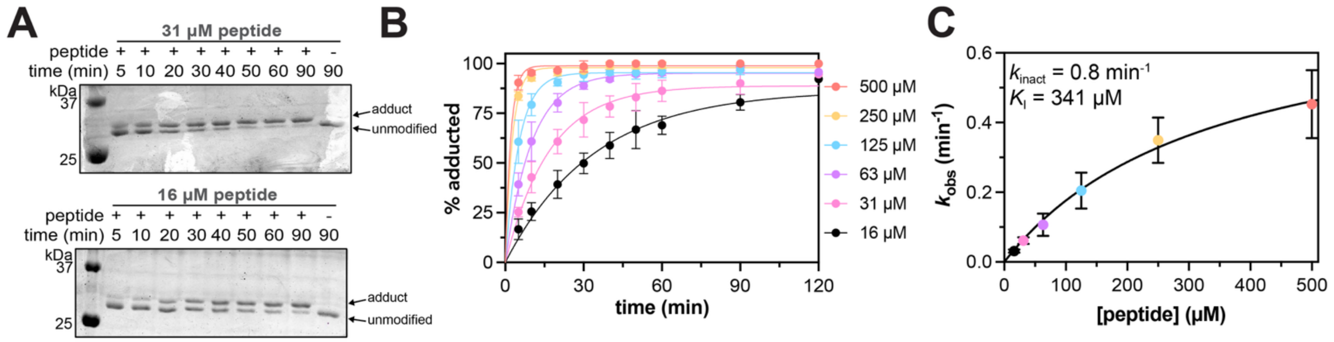


**Figure S5. Determination of $k_{inact}$ and $K_I$ parameters for Src with peptide 12.** (**A**) Representative gel-shift assays showing kinetics Src adduct formation at two different peptide concentrations. (**B**) Quantification of three replicates of gel-shift assays as a function of peptide concentration, fit to a first-order rate equation to extract $k_{obs}$ for adduct formation. (**C**) Plot of $k_{obs}$ versus peptide concentration, to extract $k_{inact}$ and $K_I$.

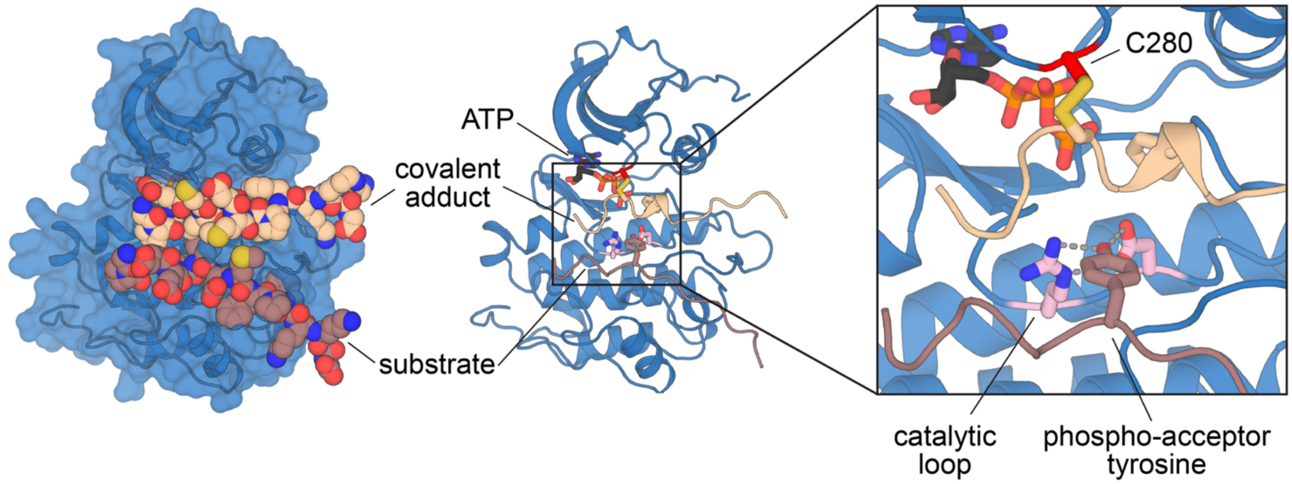


**Figure S6. AlphaFold 3 model of two peptides bound in the Src active site.** $Src_{KD}$ is modeled with a canonical, active-conformation phosphorylated activation loop. The covalent peptide (beige, modeled based on the sequence of **peptide 1**), forms a disulfide with between the -2 cysteine on the peptide and C280 on the P-loop. The substrate peptide (brown, modeled based on **peptide 1** with a C-2E substitution), shows intimate hydrogen bonding with key aspartate and arginine residues on the catalytic loop.

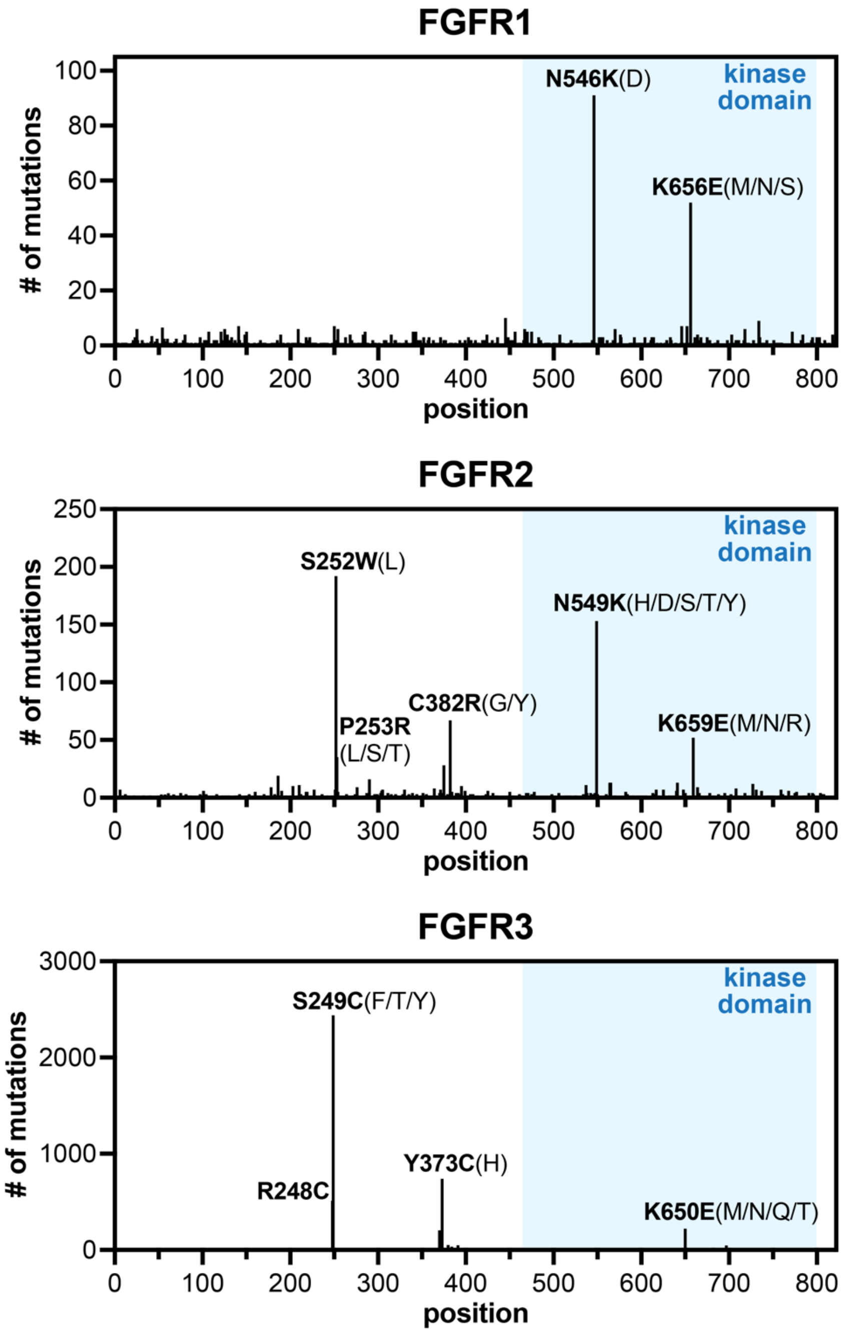


**Figure S7. The landscape of cancer mutations in FGFR kinases.** Mutational frequencies for FGFR1, FGFR2, and FGFR3 were extracted from the COSMIC database. Note that FGFR4 was omitted because it does not have a similar mutational signature in the kinase domain (blue shaded region). For prevalent mutation sites, the major substitution is shown in bold, and minor substitutions are shown in parentheses.

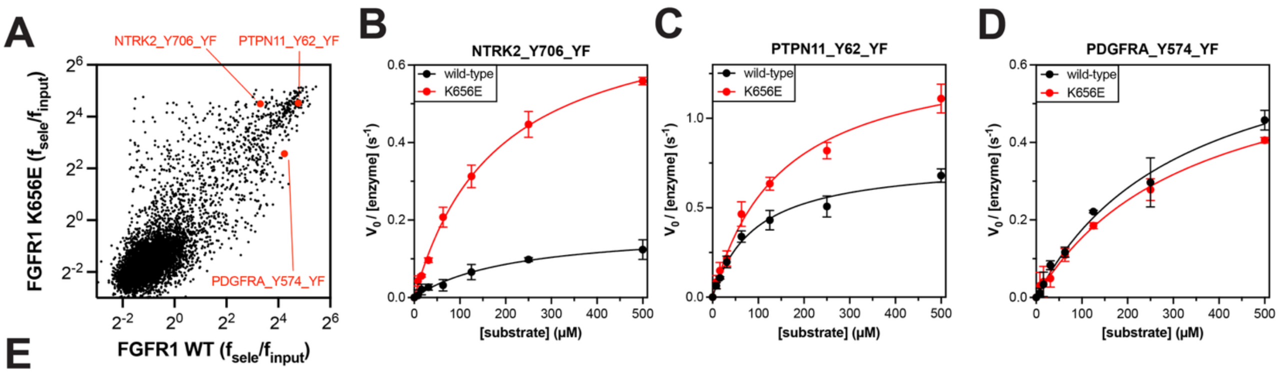


**E**

| | | FGFR1 wild-type | | | FGFR1 K656E | | |
|---|---|---|---|---|---|---|---|
| **name** | **sequence** | $k_{cat}$ ($s^{-1}$) | $K_M$ (μM) | $k_{cat}/K_M$ $(M*s)^{-1}$ | $k_{cat}$ ($s^{-1}$) | $K_M$ (μM) | $k_{cat}/K_M$ $(M*s)^{-1}$ |
| NTRK2_Y706_YF | Ac-GVFSTDYFRVGGKKKG-NH2 | 0.18 ± 0.03 | 235 ± 78 | 877 ± 315 | 0.76 ± 0.01 | 181 ± 12 | 4,247 ± 318 |
| PTPN11_Y62_YF | Ac-GQNTGDYFDLFGKKKG-NH2 | 0.76 ± 0.06 | 92 ± 12 | 8,356 ± 402 | 1.40 ± 0.06 | 149 ± 14 | 9,433 ± 850 |
| PDGFRA_Y574_YF | Ac-GGHEFIYVDPMQKKKG-NH2 | 0.75 ± 0.06 | 335 ± 64 | 2,300 ± 386 | 0.70 ± 0.11 | 364 ± 128 | 2,034 ± 348 |
| consensus, base | Ac-GPMDGDYWWLPIKKKG-NH2 | 0.88 ± 0.01 | 36 ± 1 | 24,400 ± 1,013 | 1.70 ± 0.04 | 16 ± 2 | 39,959 ± 944 |
| consensus, L+3R | Ac-GPMDGDYWWRPIKKKG-NH2 | 0.27 ± 0.01 | 155 ± 7 | 1,741 ± 119 | 1.38 ± 0.07 | 42 ± 2 | 85,414 ± 7,243 |

**Figure S8. Validation of changes in FGFR1 substrate specificity caused by the K656E mutation.** (**A**) Correlation plot (same as main figure), but highlighting validation peptides. Michaelis-Menten plots for (**B**) NTRK2_Y706_YF (mutant-selective), (**C**) PTPN11_Y62_YF (similar for wild-type and mutant), and (**D**) PDGFRA_Y574_YF. (**E**) Michaelis-Menten parameters for all FGFR1 validation peptides (including consensus peptides). Note that the intrinsic catalytic activity of the FGFR1 K656E mutant is higher than that of FGFR1 wild-type. This is corrected for in the peptide screens, but in the *in vitro* catalytic activity assays, this bias toward the mutant can be seen. Thus, a peptide that is favored by both kinases, such as PTPN11_Y62_YF, shows slightly higher activity against the mutant, and wild-type favored peptide such as PDGFRA_Y574_YF only shows a marginal preference for wild-type.

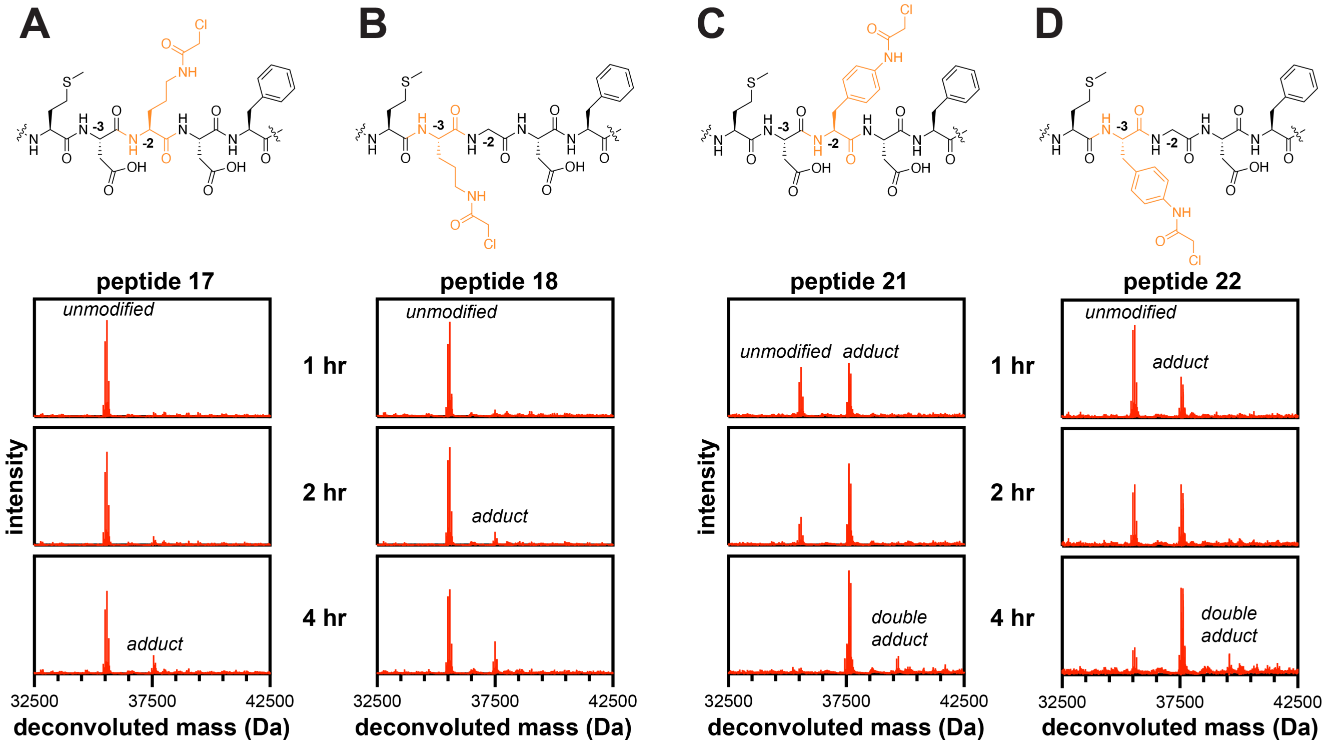


**Figure S9. Analysis of electrophile linker and position for FGFR1$_{KD}$ K656E targeting.** Structures of the -4 to 0 positions of peptides (*top*) and adduct formation mass spectra with FGFR1$_{KD}$ K656E and 100 µM peptide (*bottom*) for (**A**) **peptide 17** with a -2 electrophile and ornithine linker, (**B**) **peptide 18** with a -3 electrophile and ornithine linker, (**C**) **peptide 21** with a -2 electrophile and phenyl linker, (**D**) **peptide 22** with a -3 electrophile and phenyl linker,
